# Quantifying fluorescent glycan uptake to elucidate strain-level variability in foraging behaviors of rumen bacteria

**DOI:** 10.1101/2020.06.08.140780

**Authors:** Leeann Klassen, Greta Reintjes, Jeffrey P. Tingley, Darryl R. Jones, Jan-Hendrik Hehemann, Adam D. Smith, Timothy D. Schwinghamer, Carol Arnosti, Long Jin, Trevor W. Alexander, Carolyn Amundsen, Dallas Thomas, Rudolf Amann, Tim A. McAllister, D. Wade Abbott

**Affiliations:** Lethbridge Research and Development Centre, Agriculture and Agri-Food Canada, 5403-1st Avenue South, Lethbridge, Alberta, Canada, T1J 4B1; Department of Biological Sciences, University of Lethbridge, Lethbridge, Alberta, Canada, T1K 6T5; Max Planck Institute for Marine Microbiology, Bremen, Germany, 28359; Center for Marine Environmental Sciences, University of Bremen (MARUM), Bremen, Germany, 28359; Department of Marine Sciences, University of North Carolina, Chapel Hill, NC, United States

**Keywords:** microbiome, carbohydrate, rumen, glycoside hydrolase, fluorescent polysaccharides, yeast mannan, *Bacteroides*

## Abstract

Gut microbiomes have vast catabolic potential and are essential to host health and nutrition. An in-depth understanding of the metabolic pathways in these ecosystems will enable us to design treatments (i.e. prebiotics) that influence microbiome structure and enhance host physiology. Currently, the investigation of metabolic pathways relies on inferences derived from metagenomics or *in vitro* cultivations, however, novel approaches targeting specific cell physiologies can illuminate the functional potential encoded within microbial (meta)genomes to accurately assess metabolic abilities. Here, we present a multi-faceted study using complimentary next-generation physiology and ‘omics’ approaches to characterize the microbial adaptation to a prebiotic in the rumen ecosystem. Using fluorescently labeled polysaccharides, we identified bacteria that actively metabolize a glycan prebiotic in the rumen microbiome *ex vivo*. Subsequently, we characterized strain-level variability in carbohydrate utilization systems and predict metabolic strategies of isolated bovine-adapted strains of *Bacteroides thetaiotaomicron* using comparative whole genome sequencing, RNA-Seq, and carbohydrate-active enzyme fingerprinting.

## Introduction

Ruminants have evolved foregut digestive systems specialized in the bioconversion of recalcitrant, complex carbohydrates into energy. These catabolic processes rely on a core bacterial community composed predominantly of the genera *Prevotella, Butyrivibrio, Fibrobacter*, and *Ruminococcus*, families *Lachnospiraceae* and *Ruminococcaceae*, and orders *Bacteroidales* and *Clostridiales* (Henderson et al., 2015; Weimer, 2015). This complex microbiome is estimated to contain 69,000 carbohydrate-active enzyme (CAZyme) genes (Stewart et al., 2018) that encode extensive catalytic activities. Despite this vast sequence space and catalytic potential, the microbial conversion of plant cell walls in the rumen is suboptimal (Huws et al., 2018; Lima et al., 2019), with only 10 - 35% of consumed carbohydrates being converted into meat and milk (Varga & Kolver, 1997). Consequently, improving the efficiency of digestion with direct fed microorganisms, such as *Bacteroides* spp., or prebiotic carbohydrates that reprogram the rumen microbiome for enhanced feed conversion may help address the emerging challenges associated with sustainable production of food animals (Uyeno, Shigemori, & Shimosato, 2015).

*Bacteroides* spp. improve host digestion because they encode highly specialized carbohydrate metabolic systems called polysaccharide utilization loci (PULs) (Foley, Cockburn, & Koropatkin, 2016). The first described PUL was the starch utilization system of *Bacteroides thetaiotaomicron* (*Bt*VPI-5482) (Anderson & Salyers, 1989), and since its description, PULs that metabolize glycans with unique chemistries have been found in diverse ecosystems (Grondin, Tamura, Dejean, Abbott, & Brumer, 2017; Kruger et al., 2019; Martens, Kelly, Tauzin, & Brumer, 2014; Teeling et al., 2012). PULs are distinguished by the presence of a TonB-dependent transporter coupled to a surface glycan binding protein, known as the SusC/D-like complex, and other associated proteins that modify or bind the target glycan. These gene products function together in an orchestrated cascade to transport oligosaccharides into the periplasm where monosaccharides are released from polymeric substrates and used for primary metabolism. PULs can operate through a ‘distributive’ mechanism, which releases products (*i.e*. ‘public goods’ (Rakoff-Nahoum, Coyne, & Comstock, 2014)) to the microbial community; or a ‘selfish’ mechanism (Cuskin, Lowe, et al., 2015b), which limits product loss by confining substrate depolymerization within the cell (Abbott, Martens, Gilbert, Cuskin, & Lowe, 2015; Grondin et al., 2017). Recently, PUL-prediction (Terrapon et al., 2018) and whole-PUL characterization (Cuskin, Lowe, et al., 2015a; Larsbrink et al., 2014; Martens et al., 2011; Sonnenburg et al., 2010) have become high-priority approaches for the discovery of new CAZyme families and catalytic activities at the species (Martens et al., 2011; Sonnenburg et al., 2010) and strain levels (Hehemann, Kelly, Pudlo, Martens, & Boraston, 2012; Pluvinage et al., 2018; Rogowski et al., 2015). The primary catalysts encoded within PULs are glycoside hydrolases (GHs), which cleave glycosidic bonds by acid-base catalysis (Davies & Henrissat, 1995). GHs are divided into sequence related families that display conserved folds, mechanisms, and catalytic residues. However, these features are not necessarily representative of function as many different GH families are polyspecific (Lombard, Golaconda Ramulu, Drula, Coutinho, & Henrissat, 2014).

In addition to microorganisms that improve the efficiency of digestion, prebiotic glycans, such as yeast α-mannan (YM) and its derivative oligosaccharides (*i.e*. α-mannanoligosaccharides), can provide beneficial physiological outcomes to animals (Finck et al., 2014; Miguel, 2004; Wohlt, Corcione, & Zajac, 1998). Prebiotics do this by becoming selective nutrients for symbiotic gut bacteria, such as *Bacteroides* spp. The digestion of YM requires a collection of CAZymes targeting distinct linkages using different modes of activity, including α-mannanases and α-mannosidases (Abbott et al., 2015). CAZymes that possess these activities are commonly found in family GH38, GH76, GH92, GH99, and GH125 (Cuskin, Lowe, et al., 2015a; Gregg et al., 2011; Zhu et al., 2010); and correspondingly, these enzymes are present in PULs that target YM (*i.e*. MAN-PULs).

YM specific CAZymes and PULs are widely distributed in *Bacteroidetes*; however, individual species differ in their abilities to consume α-mannans depending on the structural complexity of the substrate (Cuskin, Lowe, et al., 2015a). For example, *Bacteroides xylanisolvens* NLAE-zl isolated from pigs reared on a diet infused with distillers’ grains could only metabolize debranched YM (Cuskin, Lowe, et al., 2015a). The pathway responsible for YM-catabolism in these strains (*i.e*., MAN-PUL1) was encoded on a transposable element, suggesting that aspects of YM metabolism can be exchanged between strains (Cuskin, Lowe, et al., 2015a). This finding is consistent with reports of specialized metabolic abilities being transferred to intestinal *Bacteroides* spp. from species that occupy ecologically distinct habitats (Hehemann et al., 2010; Hehemann et al., 2012; Pluvinage et al., 2018), facilitating their persistence within highly competitive ecosystems and adaption to spatially and culturally diversified diets.

Although major advances have been made in understanding the diversity of metabolic potential in symbiotic bacteria and the mechanisms of prebiotic utilization, establishing stabile engineered microbiomes in complex ecosystems, such as the rumen, will require more detailed knowledge of the competitive and complementary processes that drive metabolic phenotypes at the strain level. To achieve this, “next-generation physiology” (Hatzenpichler, Krukenberg, Spietz, & Jay, 2020) based approaches that identify metabolic potentials of individual bacteria, thereby providing critical insights of cellular functions and assigning cellular phenotypes, must be developed. One such approach is the application of fluorescently labeled polysaccharides (FLA-PS). FLA-PS were initially developed to demonstrate selfish uptake of marine polysaccharides in marine *Bacteroidetes* (Reintjes, Arnosti, Fuchs, & Amann, 2017), and have also been applied to the gut bacterium *Bt*VPI-5482 to confirm that YM metabolism also occurs through a selfish mechanism (Cuskin, Lowe, et al., 2015a).

Here, for the first time, we apply FLA-PS as a next-generation physiology approach to directly visualize YM metabolism by single cells in a complex rumen community and subsequently classify populations of cells using FISH. We combine this analysis with a multi-tiered study of the evolution and function of YM metabolism in bovine-adapted *B. theta* strains (*Bt*^Bov^), which adopt one of two dichotomous growth phenotypes, referred to as “High Grower” (HG) or “Medium Grower” (MG), based on the optical density of cultures after 24 hrs. Using genomics, transcriptomics, and CAZyme fingerprinting, multiple MAN-PUL architectures were identified in this study that are consistent with reports for human-associated *Bt*VPI-5482 (Cuskin, Lowe, et al., 2015a) and key differences in the YM utilization systems between MGs and HGs were uncovered. To define the mechanisms that contribute to these growth phenotypes, we present a new quantitative application of FLA-PS, which we believe has far-reaching implications for elucidating differences in substrate utilization of individual cells within complex microbial communities.

## Results

### *Ex vivo* visualization of YM-metabolising taxa within the rumen community

To assess the capability of rumen microbiota to metabolize YM, extracted rumen samples were incubated with FLA-YM and visualized on food particles and in solution (Fig. 1a,b). The total cell density in 100 μm pre-filtered rumen fluid, as determined by enumerating DAPI-stained cells, was 2.98 x 10^8^ ± 6.02 x 10^7^ cells ml^-1^ (Fig. 1c). In these complex communities, on average 6.1% ± 0.5% of cells showed uptake of FLA-YM (0% after 15 min, 6% after 3 hrs, 7% after 1 day, 6% after 3 days). Fluorescence *in situ* hybridization (FISH) using the CF968 probe (Acinas et al., 2015) specific for the phylum *Bacteroidetes* showed that 2.9 ± 0.5% of the cells showing FLA-YM uptake were members of the *Bacteroidetes*. In total, *Bacteroidetes* made up 34.8% ± 6.8% of the rumen bacterial community and only a fraction of these (~3%) showed uptake of FLA-YM (Fig. 1c). The microbial community composition of these rumen samples was determined by 16S rDNA metagenomics sequencing. The community was dominated by *Bacteroidetes*, specifically the genus *Prevotella* 1, which demonstrated that YM metabolism has penetrated distantly related members of the phylum (Fig. 1d).

**Figure 1.**
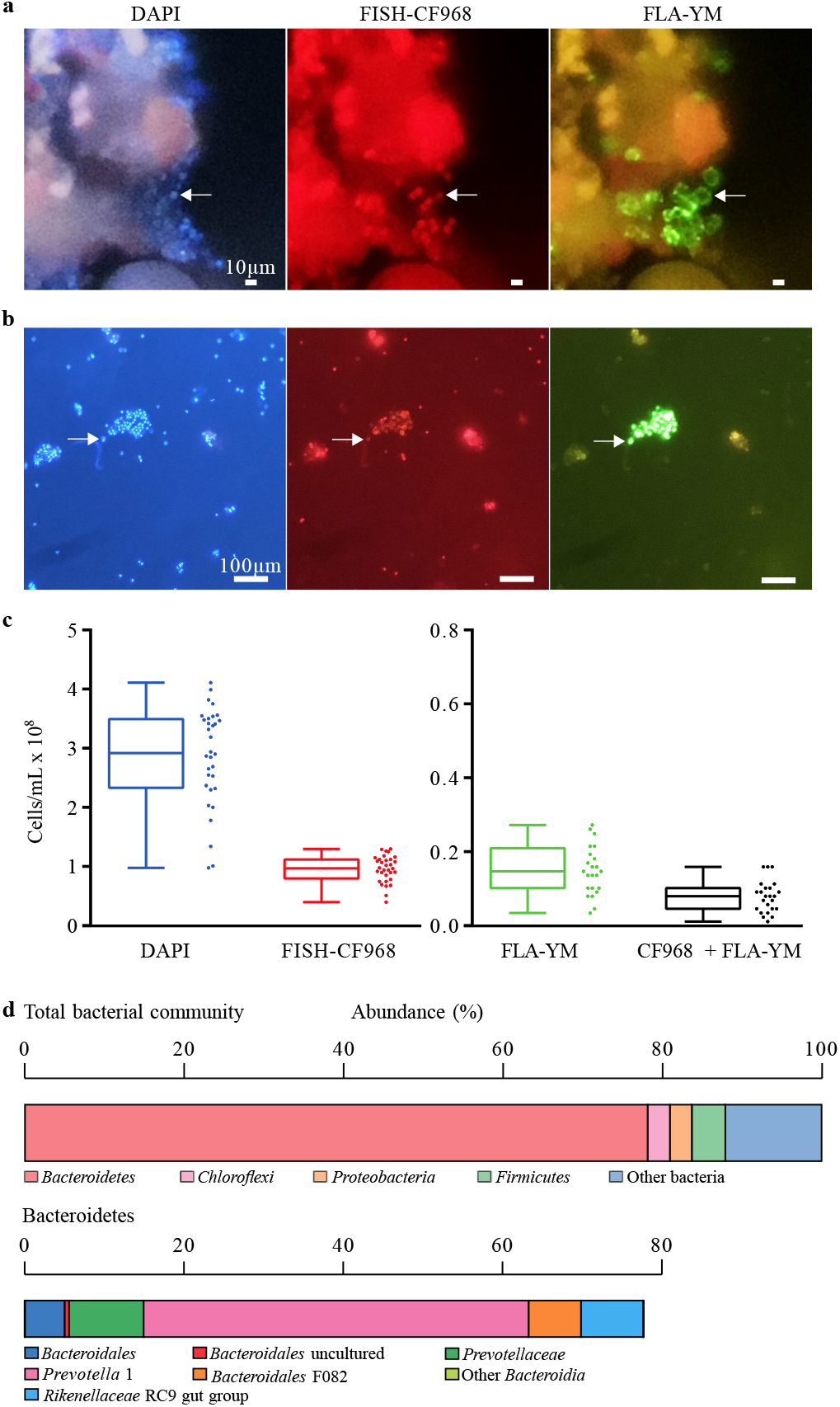
YM utilization by Bt^Bov^ isolates. Images of **(a)** 100 μm on food particles and **(b)** 10 μm filtered rumen extract incubated with FLA-YM and stained with DAPI and FISH-probe CF968. Cells were visualized by epifluorescence microscopy. **(c)** Counts of cells from FLA-YM incubated rumen extract stained with DAPI, *Bacteroides*-FISH probe (FISH-CF968), FLA-YM, and both FISH-CF968 and FLA-YM. Mean ± standard deviation shown. **(d)** 16S rRNA metagenomics sequencing data of extracted rumen communities showing most prevalent phyla (left) and the distribution of *Bacteroidetes* spp. (right). N = 4.

### Isolation and growth profiling of YM-utilizing *Bt*^Bov^ strains

Targeted isolation approaches were performed to selectively isolate bovine-adapted-bacteria that utilize YM from enriched rumen and fecal communities. Single colonies were observed within 24 hrs, with new colonies forming up to 96 hrs. In total, 50 bacterial isolates were collected and each mannan-degrading (MD) isolate was assigned a reference number (*e.g*., isolate #8 = MD8). The majority of these isolates were identified by 16S rDNA gene sequencing as strains of *B. theta* using the NCBI BLASTN database(Information, 2019), and referred to as *Bt*^Bov^ (Fig. 2a, Supplementary Table 1).

**Figure 2.**
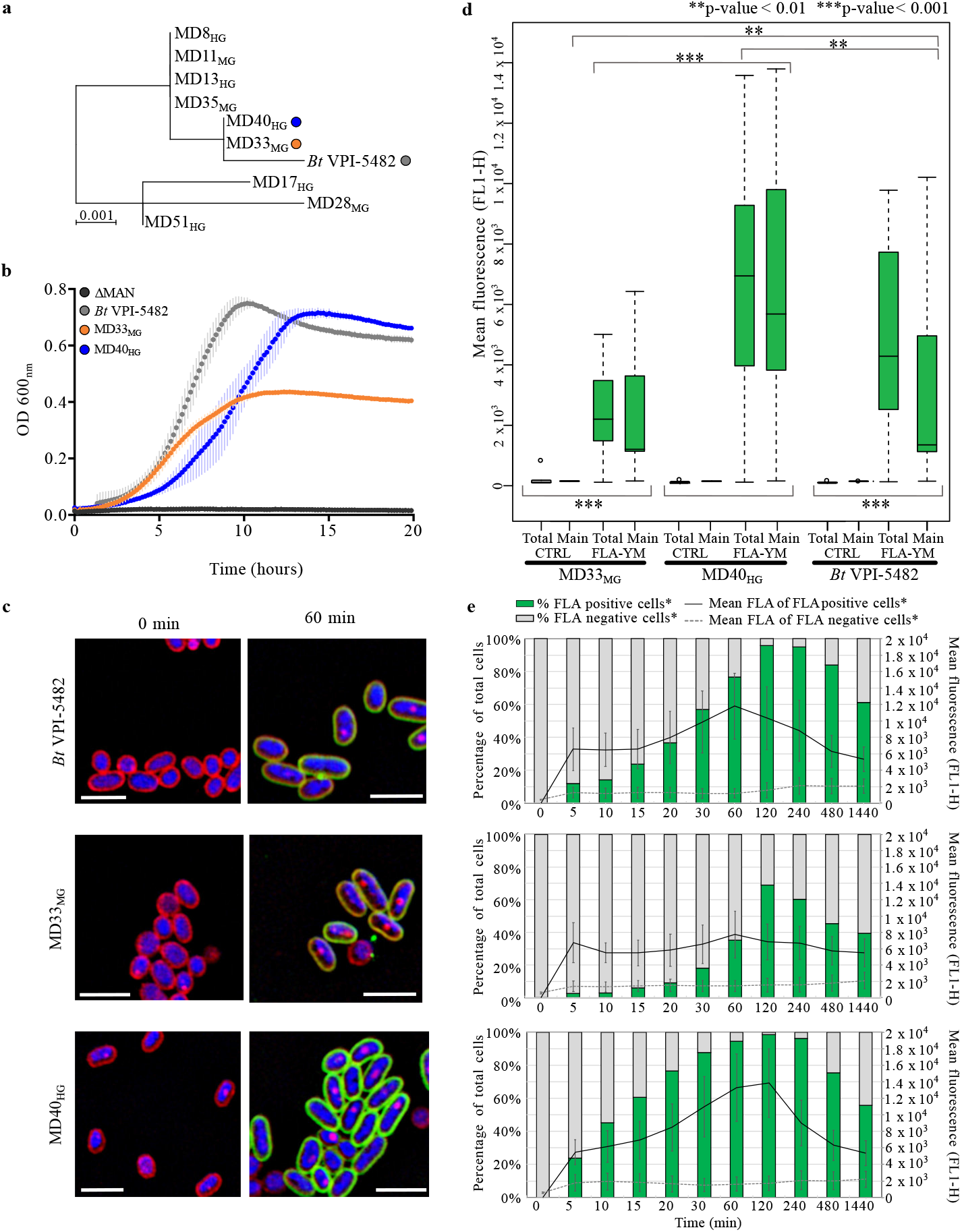
Characterization of *Bt*^Bov^ isolates. **(a)** 16S rRNA gene sequence comparison of *Bt*VPI-5482 and *Bt*^Bov^ strains. Scale represents number of nucleotide changes across horizontal axis. **(b)** Growth profiles of *Bt*VPI-5482, *Bt*ΔMAN-PUL1/2/3, MD33MG and MD4040 grown on 0.5% YM-MM. **(c)** SR-SIM of *B. theta* strains 0 min and 60 min post-incubation with FLA-YM. Cells costained with Nile Red and DAPI. **(d)** Mean fluorescence of bacterial cells cultured in FLA-YM or YM-MM (control). ** p-value < 0.01; *** p-value < 0.001; ns p-value > 0.05. (e) Bars represent percentage of total cells showing an uptake of FLA-YM in cultures sampled over time. Green = increased FLA-YM uptake; grey = no uptake. Mean cell fluorescence represented by line graph; solid line = mean of cells with FLA-YM signal; dotted line = mean of cells showing no FLA-YM uptake.

YM metabolism was confirmed for each MD strain by growth in liquid cultures using YM as the sole carbon source (Fig. 2b, Supplementary Fig. 1). Interestingly, based on their growth on *S. cerevisiae* YM, the *Bt*^Bov^ isolates and *Bt*VPI-5482 control strain were divided into two populations (Supplementary Table 2): “Medium Growers” (MGs, plateaued growth at OD_600_~0.4 after 24 hrs) and “High Growers” (HGs, plateaued growth at OD_600_~0.7 after 24 hrs). Notably, the 16S rDNA gene topology of the *Bt*^Bov^ isolates did not reveal a discernable relationship with the growth phenotype (Fig. 2a).

### Visualization and quantification of differential FLA-YM uptake by *Bt*^Bov^ isolates

To determine if the rate of glycan uptake varied between the two growth types, representative strains were incubated with FLA-YM. *Bt*VPI-5482, MD33_MG_, and MD40_HG_ cells became fluorescent, whereas cells incubated with unlabeled YM did not (Fig. 2c). Phenotypic differences in FLA-YM uptake over time were determined by splitting cell populations by flow cytometric gating into FLA-positive and FLA-negative cells (Supplementary Fig. 2a). The total fluorescence intensity (rate of uptake) was significantly different between the growth phenotypes (MD40_HG_ vs. MD33_MG_ *t*(10) = 4.4, *P*-value = 0.001; MD40_HG_ vs. *Bt*VPI-5482 *t*(10) = 3.9, *P*-value = 0.003; *Bt*VPI-5482 vs. MD33_MG_ *t*(10) = 4.0, *P*-value = 0.002) (Fig. 2d). The change in mean fluorescence of the three strains showed a similar temporal pattern, increasing from 0 to 120 min, peaking at 120 min, and declining from 120 to 1440 min (Fig. 2e). Although all three strains showed uptake of FLA-YM after 60 min, MD33_MG_ had the lowest fluorescence intensity at each time point. MD40_HG_ displayed the highest fluorescence intensity, 2.7-fold higher than MD33_MG_. *Bt*VPI-5482 displayed a fluorescence intensity between the two bovine strains for each time point, with a peak value 1.9-fold higher than MD33_MG_. Additionally, MD40_HG_ cells showed a more rapid uptake (22% at 5 min), *Bt*VPI-5482 cells showed an intermediate rate of uptake (9% at 5 min), and MD33_MG_ had the slowest uptake rate (2% at 5 min) (Fig. 2e, Supplementary Fig. 2b).

To test if the phenotypic differences were inherited between generations, we measured the differences in FLA-YM uptake with and without prior exposure to YM. The cultures continued to display the same phenotypic uptake patterns (MD40_HG_ and *Bt*VPI-5482 higher uptake, MD33_MG_ low uptake, Fig. 3a-c). However, previous exposure to YM resulted in a heightened cellular response as indicated by more rapid rates of FLA-YM uptake relative to cultures previously grown on mannose-MM (Fig. 3b-d). All cultures grown on mannose-MM reached a lower mean fluorescence and the temporal change in mean fluorescence was slower, with quantifiable uptake occurring only after 4 to 8 hrs.

**Figure 3.**
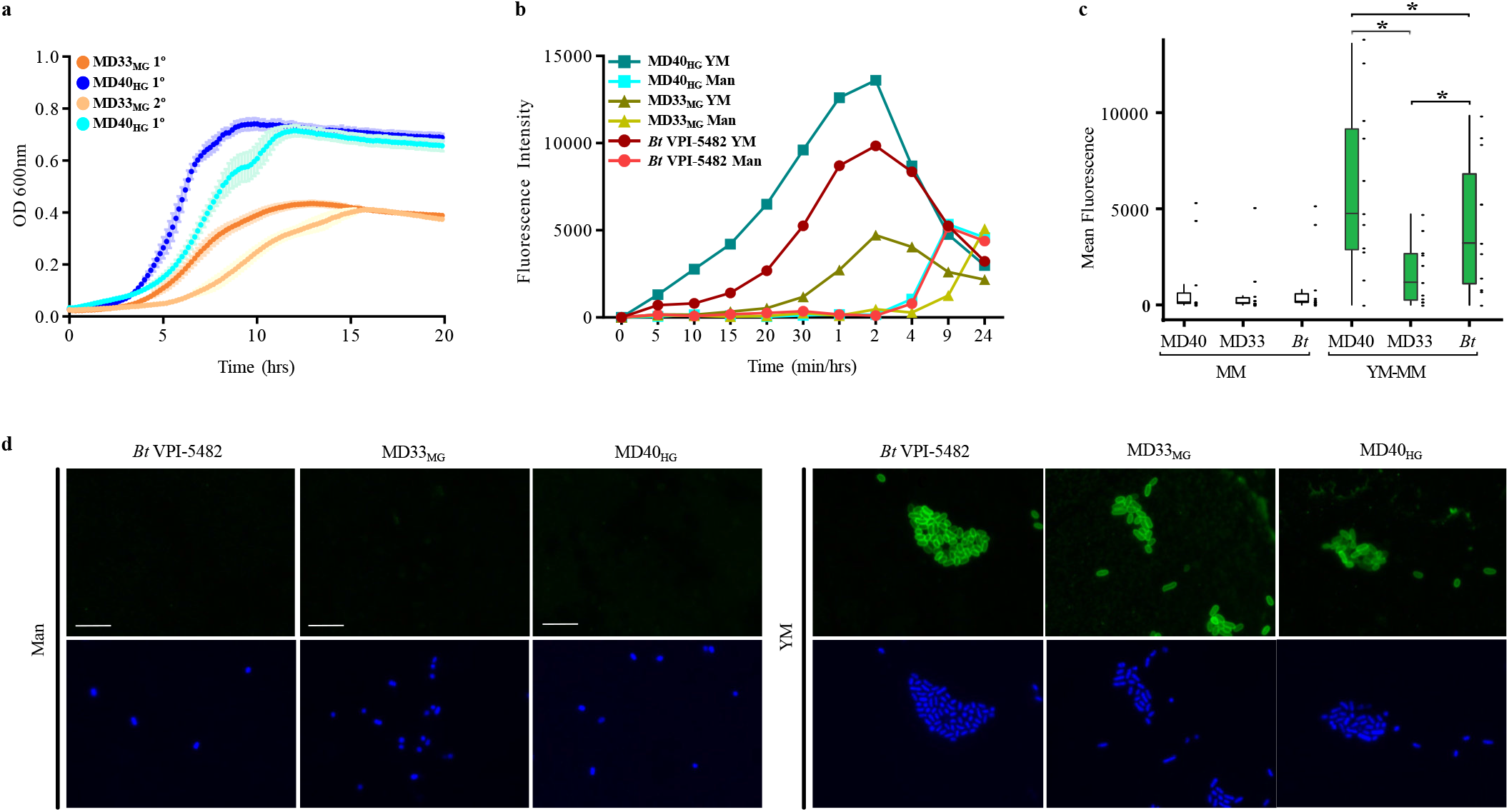
Reproducible YM foraging behaviors in *Bt*^Bov^ isolates. **(a)** Growth profiles of MD33_MG_ and MD40_HG_ inoculated from cultures cultivated overnight on mannose-MM (1°) or YM-MM (2°). **(b)** Fluorescence intensity of individual cells of *Bt*^Bov^ strains incubated in FLA-YM. Cells used to inoculate FLA-YM cultures were previously grown overnight in either YM-MM (YM) or Mannose (Man) (N = 1). **(c)** Mean fluorescence of cells cultured in FLA-YM with significance (p < 0.05) between the cultures previously grown on YM (right) indicated by asterisk (N = 10,000). **(d)** SR-SIM images of *B. theta*, MD33_MG_, and MD40_HG_ cultured in Man or YM and incubated with FLA-YM.

### Characterization of genotypes by PUL delineation

Whole genome sequencing and *de novo* assembly were used to identify genes involved in YM metabolism. SPAdes (Bankevich et al., 2012) assembly output and average nucleotide identity based on BLAST+ (ANIb) (GmbH, 2019; Richter, Rossello-Mora, Oliver Glockner, & Peplies, 2016) are shown in Supplementary Table 2. The ANIb results supported the 16S rDNA gene sequence data, confirming that each isolate was a strain of *B. theta*. Further, comparative genomics revealed that these strains have acquired unique CAZome repositories (Fig. 4a) and PUL updates (Supplementary Fig. 3). features that may assist with their colonization of the bovine gut and represents opportunities for developing bovine-adapted probiotics (Supplementary Discussion).

**Figure 4.**
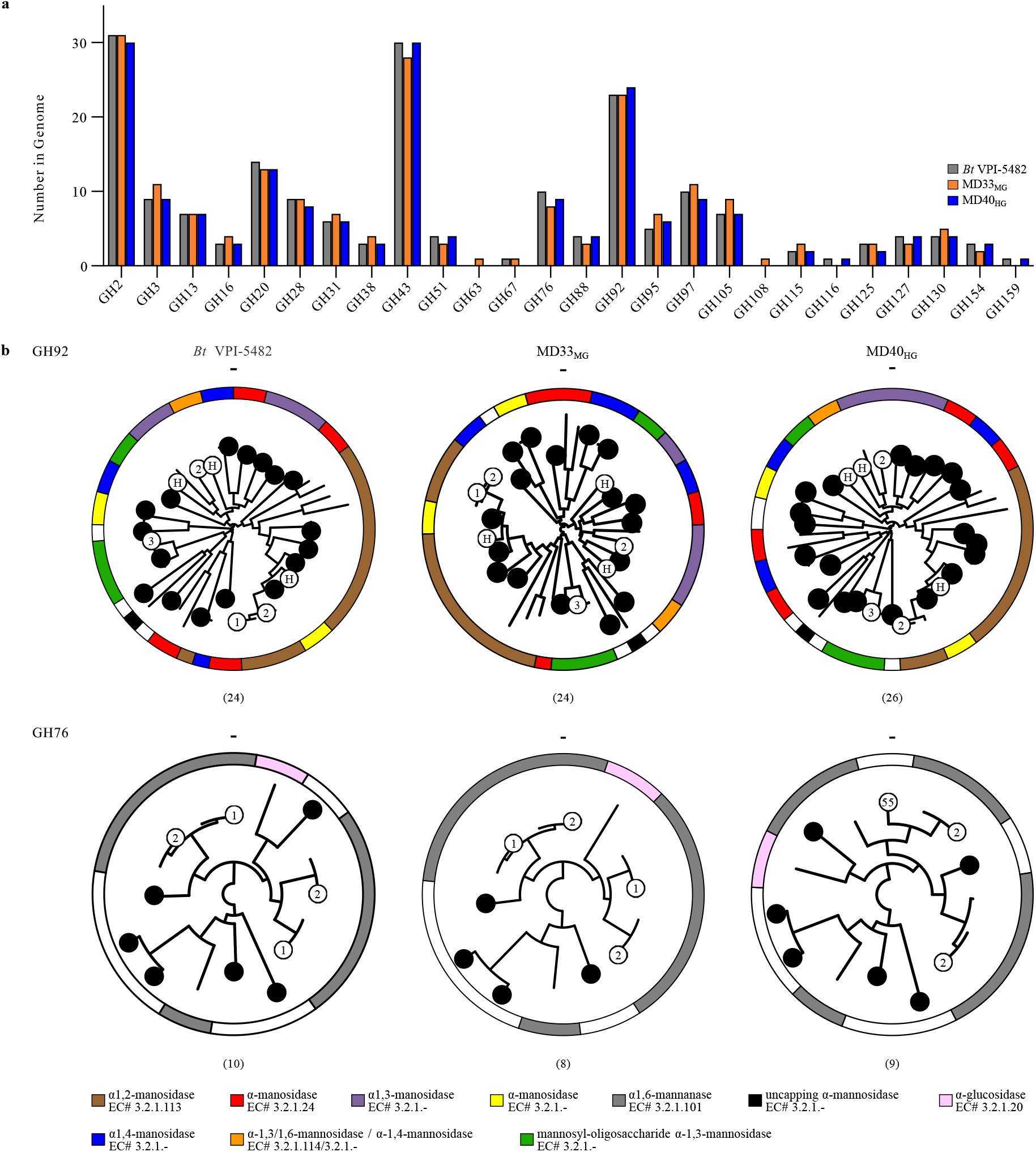
CAZyme fingerprinting of YM metabolism by *Bt*^Bov^ isolates. **(a)** GH enzyme families encoded within the genomes of MD40_HG_ and MD33_MG_ that differ in total number of sequences. *B. theta* sequences are provided as a reference for each GH family. **(b)** Phylogenetic trees of characterized GH92s and GH76s, and *Bt*^Bov^ sequences generated with SACCHARIS(Jones et al., 2018). Circles represent *Bt*^Bov^ sequences and white circles highlight sequences from PULs with mannan activity: 1, 2, or 3 = MAN-PUL 1, 2, or 3, respectively; H = HMNG-PUL; 55 = BtMD40 PUL55. Activities assigned to characterized enzymes within each clade for GH92 and GH76 are depicted using the provided legend. Outer ring represents characterized specificities. Numbers in parenthesis indicate the total number of enzymes within each strain.

Reconstruction of the three YM-specific PULs and alignment with *Bt*VPI-5482 MAN-PULs determined that there was a high level of synteny among all strains in these pathways (Supplementary Fig. 4a). MAN-PUL2 and 3 were absolutely conserved; whereas, MAN-PUL1, a PUL tailored for the consumption of mannan from *Schizosaccharomyces pombe*(Cuskin, Lowe, et al., 2015a) (Supplementary Fig. 1b), was only present in *Bt*VPI-5482, MD33_MG_, and MD35_MG_. The presence of MAN-PUL1 in two MGs indicated this pathway was not responsible for the HG phenotype. The HMNG-PUL, which is specific for digestion of high mannose N-glycans and not activated by YM in *B. theta* (Cuskin, Lowe, et al., 2015a), was also conserved in each of the *Bt*^Bov^ genomes.

### CAZome fingerprinting

To determine if there was amino acid sequence divergence within MAN-PULs, and potentially the function of homologous enzymes, polyspecific CAZyme families GH92 and GH76 were analyzed by SACCHARIS (Jones et al., 2018). Enzyme sequences from GH92 and GH76 were embedded into phylogenetic trees comprised of all characterized enzyme sequences from each family (Supplementary Fig. 4b,c). Notably, every sequence within MAN-PUL1, 2, and 3 displayed the highest level of amino acid sequence conservation with its syntenic homolog. This suggested that each PUL is under strong selective pressure to function as an intact catabolic system. To determine if CAZyme sequences were conserved in other potential α-mannan degrading PULs, a genome-wide approach (*i.e*., CAZome fingerprinting) was used (Jones et al., 2018). Each isolate encoded between twenty-four and twenty-six GH92s, and eight or nine GH76s (Fig. 4b, Supplementary Fig. 4b,c). Only MD17_HG_ and MD51_HG_ displayed identical conservation for GH76; whereas, every GH92 tree was unique.

Topological differences were observed for other α-mannan active enzyme families (e.g. GH38, GH99, and GH125), suggesting that despite the high level of functional conservation within the MAN-PULs, metabolic specialization in α-mannan consumption between these strains may be encoded within orphan PULs (Ndeh et al., 2017). Therefore, the contributions of two exogenous GH76s to the foraging behavior of HGs and MGs was investigated. BtGH76-MD40 is a surface-exposed GH76 inserted into PUL55 of HGs (Fig. 4b) and is active on intact *S. cerevisiae* and *S. pombe* YM (Jones et al., 2020). BT_3782 is a periplasmic endo-α-mannanase that generates small oligosaccharide products (Cuskin, Lowe, et al., 2015b). Addition of recombinant BtGH76-MD and BT_3782 to pure cultures of MD33_MG_ did not augment the MG growth phenotype (Supplementary Fig. 5), suggesting that acquisition of BtGH76-MD40 or augmented endo-mannanase activity were not responsible for the HG phenotype.

### Differences in YM import between phenotypes

The differential transport kinetics of FLA-YM (Fig. 2c-e) and absence of genetic differences in PUL structure between phenotypes (Supplementary Fig. 4a) suggested that glycan transport processes may be responsible for the MG and HG growth phenotypes. Alignment of the SusC-like amino acid sequences from MAN-PUL 1, 2, and 3 and HMNG-PUL from *Bt*VPI-5482, MD33_MG_ and MD40_HG_ revealed that proteins cluster into functional clades (Fig. 5a). Furthermore, when the SusC-like and SusD-like transport proteins from each MD strain were aligned with BT_3788 (Fig. 5b) and BT_3789 (Fig. 5c) of *Bt*VPI-5482, respectively, the proteins partitioned exclusively into clades associated with either the MG or HG phenotype. This result is in contrast with the 16S (Fig. 2a) and whole-genome (Supplementary Table 2) alignments, which showed no correlation with growth profiles. Interestingly, this pattern does not exist for MAN-PUL1 or MAN-PUL3 as the SusC-like proteins in these pathways are highly conserved (Fig. 5a), suggesting that syntenic conservation may not always reflect sequence-function relationships.

**Figure 5.**
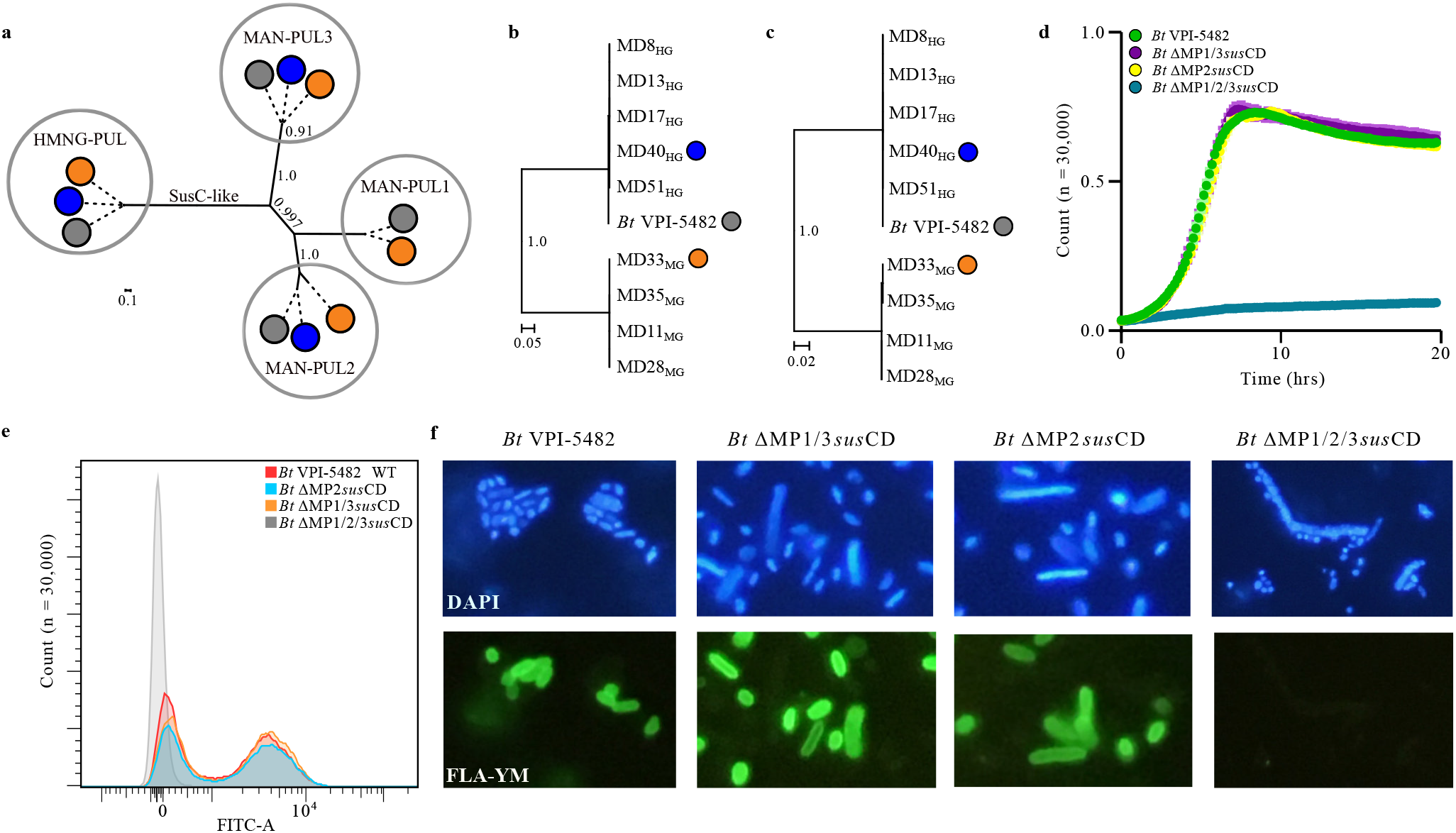
Divergence of MAN-PUL SusC/D-like proteins. **(a)** Phylogenetic tree of all the SusC-like MAN-PUL protein sequences from BtVPI-5482 (grey), MD33_MG_ (orange), and MD40_HG_ (blue). Phylogenetic trees of **(b)** SusC-like and **(c)** SusD-like amino acid sequences encoded in MAN-PUL2 of each *Bt*^Bov^ strain. Bootstrap values above 70% are indicated at branch points. Scale represents number of amino acid substitutions per site. **(d)** Growth of *Bt*VPI-5482 wild-type and MAN-PUL *susC/D*-like mutants on 0.5% YM-MM. **(e)** Enumeration (N = 30,000) and **(f)** epifluorescence microscopy images of the uptake of FLA-YM by *Bt*VPI-5482 wild-type and mutant strains after 2 hours incubation. Cells counterstained with DAPI (blue).

To study how transporters affect YM uptake, different combinations of *susC/D*-like genes were excised from the *Bt*VPI-5482 MAN-PULs. Three mutant strains were produced: a MAN-PUL2 *susC*-like and *susD-*like gene knock-out strain (ΔMP2*susCD*), a MAN-PUL1 & 3 *susC*-like and *susD*-like deletion mutant (ΔMP1/3*susCD*), and a strain with all three sets of *susC*-like and *susD*-like genes deleted (ΔMP1/2/3*susCD*). Because MAN-PUL1 is absent in every HG except *Bt*VPI-5482, we can conclude it has no effect on YM transport efficiency and that the ΔMP1/3*susCD* mutant essentially operates as a *Bt*^Bov^ MAN-PUL3 *susCD* knock-out strain. The mutants, along with *Bt*VPI-5482, were grown on YM-MM to assess how the loss of transport complexes impacted growth on YM (Fig. 5d). Surprisingly, the mutants retained an identical growth profile to the wild-type, with the exception of the triple knock-out mutant (ΔMP1/2/3*susCD*), which displayed no growth. Furthermore, when the mutants were incubated with FLA-YM, to study the impact on uptake rates, they displayed identical rates to the wild-type, with only the triple deletion mutant having a complete loss of FLA-YM import (Fig. 5e,f). These results demonstrated that the SusC-like/SusD-like proteins from MAN-PUL2 and 3 in *Bt*VPI5482 are functionally redundant. This observation is in contrast with the differential transport of RG-II substrates purified from wine and apple pectin by a tandem set SusC-like/SusD-like systems in *Bt*VPI-5482 (Ndeh et al., 2017). Although the absence of genetic tools prevented investigating the interplay between the MD33_MG_ transporters, the sequence divergence existing between MAN-PUL2 SusC/D/E-like proteins from MD33_MG_ and MD40_HG_ (S4 Table) suggested that the dichotomous MG and HG growth phenotypes may result from differential transport through these complexes.

### Comparative analysis of gene expression between *Bt*^Bov^ growth phenotypes

RNA-seq was performed on *Bt*VPI-5482, MD33_MG_, and MD40_HG_ cultured on either mannose or YM to explore differential patterns in expression of the enzymes and transporters in the MAN-PULs and identify any distally expressed genes. MAN-PUL2 and 3 pathways were activated in all three bacteria, and MAN-PUL1 was activated in *Bt*VPI-5482 and MD33_MG_ (Fig. 6a, Supplementary Fig. 6) consistent with previous reports for *Bt*VPI-5482 (Martens et al., 2011) (see Supplementary Discussion). To confirm that gene expression was representative of protein production, a C-Myc tag was fused to the C-terminal of the MAN-PUL2 SusD-like protein (BT_3789) in the chromosome of *Bt*VPI-5482. Extracellular display of BT_3789 was demonstrated using antibodies directed at C-Myc when this bacterium was cultured on YM but not mannose (Supplementary Fig. 7).

**Figure 6.**
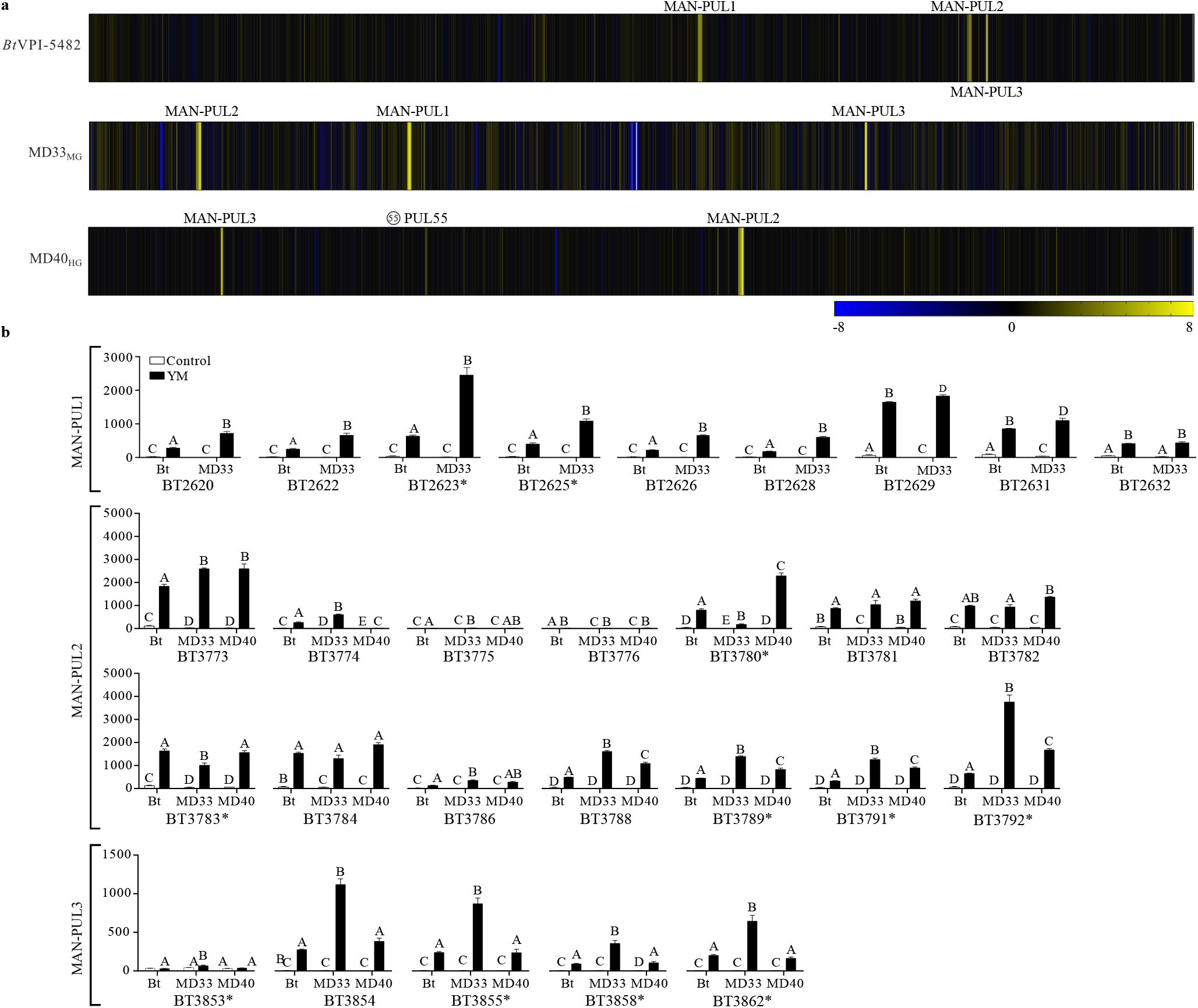
RNA-Seq analysis of *Bt*VPI-5482, MD33_MG_, and MD40_HG_ cultured in YM. **(a)** Log_2_ expression ratios of each transcript of *Bt*VPI-5482, MD33_MG_, and MD40_HG_. Length of heap map is representative of genome size (*Bt*VPI-5482 = 6.26 Mb, MD33_MG_ = 6.28 Mb, MD40_HG_ = 6.16 Mb). Location of MAN-PULs and MD40-PUL55 (white star) shown. Yellow indicates an increase in transcript expression compared to the control, while blue is decreased expression. **(b)** TPM values of *susC*-like genes from MAN-PUL1/2/3 showing if values within each gene are significantly different between the strains; different letters represent statistically significant values (*e.g*. A vs B) for each separate gene histogram; asterisk indicates SPII predicted proteins. Log_2_ fold-change compares gene expression of YM cultures normalized to mannose, N=3, p ≤ 0.01; except BT_3853 and HMNG-PUL gene expression is not significantly (p > 0.05) different between the two treatments. Significantly different (p < 0.05) TPM expression of the susC-like genes between *Bt*VPI-5482 and *Bt*^Bov^ strains; statistical comparison of other MAN-PUL genes between strains were not calculated.

The TPM values for every homologous gene transcript from MD33_MG_, MD40_HG_, and *Bt*VPI-5482, was analyzed. Surprisingly, the *sus*-like genes (BT_3788 and BT_3789) and the surface enzyme transcripts (BT_3792, BT_2623, and BT_3858) of MD33_MG_ consistently displayed significantly higher expression levels than the HG strains (Fig. 6b, Supplementary Fig. 6b). These values ranged between 6.2-log_2_ to 7.7-log_2_, suggesting that the expression level of gene products involved in outer membrane processing and intracellular transport is negatively correlated with growth proficiency. The only example of an enzyme that is expressed at a significantly higher level in the MD40_HG_ strain was BT_3780 (12.5-fold higher than MD33_MG_), which encodes a GH130 that is active on β-1,2-mannosides (Cuskin, Basle, et al., 2015). This linkage is found in *Candida albicans* cell wall, and therefore, most likely represents an off-target effect of MAN-PUL2 induction.

### Differences in YM hydrolysis and import

The enzymatic processing of YM (amount of YM products, extent of YM utilization, and total free mannose present in the post-growth supernatants) by each culture was analyzed using a combination of methods. Thin layer chromatography (TLC) (Fig. 7a) revealed that there was no detectable free mannooligosaccharides or mannose in the supernatant of the YM-MM negative control. Consistent with this observation, the *Bt*MAN-PUL1/2/3 deletion mutant (Δ*MAN-PUL1/2/3*) did not grow on YM and was unable to release products into the medium. *Bt*VPI-5482 and each of the HGs generated a similar product profile, with a noticeable loss of YM signal and faint detection of oligosaccharides and mannose. In contrast, the post-growth media of MGs contained more mannose and had residual YM (Fig. 7a). Gas chromatography-mass spectrometry determined that the quantity of total mannosides (YM and oligosaccharides) in the supernatant was 1.48 and 1.40-fold higher for MD33_MG_ (1.36 ± 0.29) than *Bt*VPI-5482 (0.92 ± 0.04) and MD40_HG_ (0.97 ± 0.05), respectively (Fig. 7b). Furthermore, post-growth *Bt*VPI-5482 and MD40_HG_ cultures, but not MD33_MG_, showed (p < 0.05) lower total mannose concentration in the media relative to the YM-MM negative control (Fig. 7b). This suggests that, consistent with their higher growth densities (Fig. 2b) and thin layer chromatography, *Bt*VPI-5482 and MD40_HG_ consume more YM.

**Figure 7.**
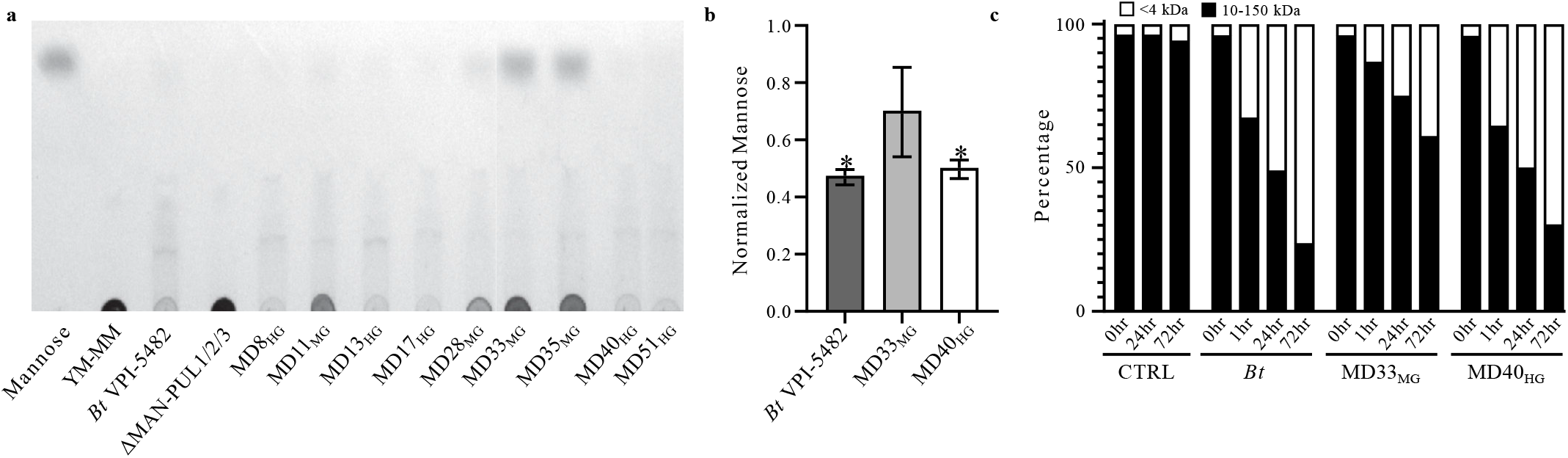
Differential transport of YM by *Bt*^Bov^ strains. **(a)** TLC analysis of post-growth supernatants (50 hrs) of MD isolates grown on 0.5% YM-MM. *B. theta* = *Bt*VPI-5482, ΔMAN = *Bt*ΔMAN-PUL1/2/3, Man = mannose standard. **(b)** Quantification of total mannose (free and polymerized) in supernatants of *Bt*VPI-5482, MD33_MG_, and MD40_HG_ grown in 0.5% YM-MM for 21 hrs. Normalized to a no cell control, N=2, asterisk indicates significant difference (p < 0.05) relative to control. **(c)** Percent distribution of FLA-YM hydrolysis products in the supernatants of *Bt*VPI-5482, MD33_MG_, and MD40_HG_ grown in 0.2% FLA-YM over time. N=3.

To determine if surface α-mannanases generate different product profiles, cultures of *Bt*VPI-5482, MD33_MG_, and MD40_HG_ were incubated with FLA-YM and the products were analyzed by fluorescence-coupled high-performance liquid chromatography. In *Bt*VPI-5482 and MD40_HG_, there was preferential hydrolysis of large FLA-YM products (> 10 kDa, ~55-mer), which was accompanied by a relative accumulation of lower molecular weight (< 4 kDa; ~22-mer) hydrolysis products (Fig. 7c, Supplementary Fig. 8).

## Discussion

The gut microbiome plays an integral role in digestion and nutrient acquisition. Improved understanding of the functional potential encoded within members of the microbiota is still required to define metabolic abilities and microbial-prebiotic interactions. Next-generation physiology approaches represent promising strategies to rapidly assign cellular phenotypes and can consolidate genomic predictions (Hatzenpichler et al., 2020). By combining phenotypic and sequencing approaches, we have conducted a high-resolution study of differential YM utilization by isolated bovine-associated bacterial strains. In liquid culture, the isolates displayed one of two growth patterns: MG or HG; trends that were independent of taxonomic relationships (Fig. 2a,b, Supplementary Table 2). This showed that closely related *Bacteroides* spp. have evolved different foraging strategies for the same substrate. FLA-PS were successfully used to visualize (Fig. 2c), and quantify the accumulation (Fig. 2d) and uptake rate (Fig. 2e) of YM products in bacterial cells, confirming that HGs use a selfish mode of metabolism on this substrate, as previously reported for *Bt*VPI-5482 (Cuskin, Lowe, et al., 2015a; Hehemann et al., 2019). In contrast, the MG strains consumed less YM and released mannose into the medium (Fig. 7a,b), suggesting that MGs are less adept at YM catabolism and display some properties consistent with distributive metabolism (Fig. 2b and 7c).

Comparative genomics revealed genotypes with high synteny across genomes and MAN-PUL pathways, with few exceptions. Perhaps the most interesting genetic anomaly is the sequence variability of the MAN-PUL2 SusC/D-like proteins, which elegantly branch into two clades coinciding with the HG and MG growth phenotype (Fig. 5b,c), as well as differential rates and total levels of FLA-PS uptake (Fig. 2c-e, Supplementary Fig. 2b). Previously, it was shown that the amino acid homology of a SusD-like protein involved in utilization of two different fructans was low between two strains of *B. theta*, despite their taxonomic similarity (Hehemann et al., 2019), highlighting that syntenic genes within PULs can evolve independently. Here we report that SusC/D-like amino acid sequences from the major PUL involved in metabolism of YM correlate with differential utilization of a common substrate (Fig. 5b,c). Because there is no perceived difference in the structures of surface enzymes encoded within the MAN-PULs (Supplementary Fig. 4), the outer surface endo-α-mannanses are expressed at lower levels in HGs (Fig. 6b), and the addition of exogenous endo-GH76s to MG growth cultures did not augment MG growth (Supplementary Fig. 5), the higher growth and faster disappearance of large YM products in MD40_HG_ cultures reflect a more efficient transport process (Fig. 7). Whether this is the direct result of higher transporter efficiency in MD40_HG_ or indirect result from impoverished transport leading to product inhibition of surface enzymes in MD33_MG_ is unclear. Intriguingly, deletion of MAN-PUL1/3 *susC/D* or the MAN-PUL2 *susC/D* did not impede growth of *Bt*VPI5482 on YM or uptake of YM (Fig. 5d-f), suggesting SusC/D-like pairs in MAN-PUL2 and 3 are functionally redundant in HGs. Based upon sequence identity (S4 Table), MGs possess one compromised SusC/D/E complex (MAN-PUL2) and one high-performing SusC/D complex (MAN-PUL3), which are regulated differently between the strains. In MD33_MG_, the MAN-PUL3 *susC-like* gene (*bt3854* homolog) is expressed at level similar to the MAN-PUL2 *susC-like* gene (*bt3788* homolog), and at a level 2.9-fold higher than its homologous gene in MD40_HG_ (Fig. 6b). Higher expression of outer surface proteins in MD33_MG_ is a consistent pattern (Fig. 6b), suggesting it is trying to compensate for its growth inefficiencies, but is unable to do so. Despite the higher expression levels of the MAN-PUL3 SusC/D-like complex in MD33_MG_, and the ability of the MAN-PUL3 SusC/D-like complex to compensate for deletion of the MAN-PUL2 transporter in *Bt*VPI-5482 (Fig. 5d,e), the MAN-PUL3 SusC/D-like complex in MD33_MG_ is unable to rescue the MG growth phenotype. Thus, the SusC/D-like complexes in MAN-PUL2 and 3 appear to compete for substrates and the inefficiencies of transport ascribed to the MAN-PUL2 complex are related to its ability to transport, but not recruit, YM substrates. Further biochemical and structural studies of the MAN-PUL2 SusC/D-like proteins are warranted to tease apart these results.

The “Nutrient Niche Hypothesis” (Freter, Brickner, Botney, Cleven, & Aranki, 1983) suggests that metabolic abilities are determined by the creation and filling of ecological nutrient niches. In theory, these relationships could be in response to the introduction of a new dietary glycan (*i.e*. prebiotic), resulting in the selection for or adaptation of a bacterium with the metabolic capacity to consume it. In this study, the MG and HG phenotypes represent a variation on this theme, as two closely related populations (> 98% identity) adapted to the colonization of a common host (Supplementary Fig. 3) display different (Fig. 2b, Supplementary Fig. 1), yet reproducible (Fig. 3a) and inducible (Fig. 3b-d), foraging behaviors on the same substrate. These findings raise several unsolved questions related to the existence and persistence of MGs, and potentially other glycan foragers that are less adept at substrate utilization, in the rumen. If HGs have a superior capacity for YM metabolism, why are MGs not eliminated by competitive exclusion? And if MGs have restricted abilities to digest YM and / or transport YM products (Fig. 7), why are these PULs not selected against and excised from the genome? The existence of multiple metabolic phenotypes suggests that ecological selection factors may be responsible. Firstly, feeding strategies, such as distributive metabolism, may foster beneficial syntrophic relationships at multiple levels within a community (Koropatkin, Cameron, & Martens, 2012; Morris, Henneberger, Huber, & Moissl-Eichinger, 2013). The generation of public goods (Rakoff-Nahoum et al., 2014) by MGs provides nutrients to species that are incapable of digesting YM. This event would increase the richness of the community, and potentially, result in the generation of additional secondary metabolites that benefit the lifestyle of MGs. Furthermore, it has been shown that both the concentrations and complexity of available substrate cause differential selection of distributive or selfish foraging strategies (Reintjes et al., 2020; Rogowski et al., 2015; Sarmento, Morana, & Gasol, 2016). Secondly, *Bacteroides* spp. are generalists with the capacity to utilize a wide variety of substrates available in the diet of their hosts and glycan responses are prioritized in *Bacteroides* spp. (Rogers et al., 2013; Tuncil et al., 2017). MGs may possess a different substrate hierarchy than HGs, and correspondingly, display more prowess for consuming chemically distinct glycans. Alternative substrate priorities would reduce the competitive burden on MGs when provided with complex diets. In this regard, the acquisition of new CAZymes or PULs that endow a microorganism with an ability to consume new substrates, has been hypothesized to occur by horizontal gene transfer and is linked to spatial and cultural dietary habits (Hehemann et al., 2010; Pluvinage et al., 2018).

Comparison of the MD40_HG_ and MD33_MG_ CAZomes confirmed that there are many GH families, encoding different enzyme specificities, that vary in number (Fig. 4a). Closer inspection of GH3 and GH16, two polyspecific GH families active on β-linkages, revealed CAZyme updates within a PUL in MD33_MG_ (Fig. 8). This suggests that the acquisition of a putative β-glucan metabolic pathway, and potentially others, may provide a colonization advantage for MD33_MG_ despite its weakened potential to metabolize YM. Recently, β-glucan utilization pathways were shown to have independently evolving genes that result in the expansion of protein specificity and glycan targets(Dejean et al., 2020). Thus, clustered mutations in PULs could unlock previously inaccessible nutrient niches. Conversely, however, there is the risk of impeding nutrient acquisition, as exhibited by the restricted β-glucan utilization of polyspecific proteins in *Bacteroides fluxus* (Dejean et al., 2020), and potentially, the inefficiencies of YM uptake governed by transporter specificity or efficiency. Further investigation of total CAZome function and transporter selectivity and efficiency encoded within genomes at the strain level will reveal how microorganisms living in partnership or competition within complex ecosystems tune their metabolic responses to complex dietary landscapes. Coupling ‘omics’ methods and functional methods, such as FLA-PS, will help usher in a new frontier for the assignment of metabolic traits to bacterial populations within microecological food webs.

**Figure 8.**
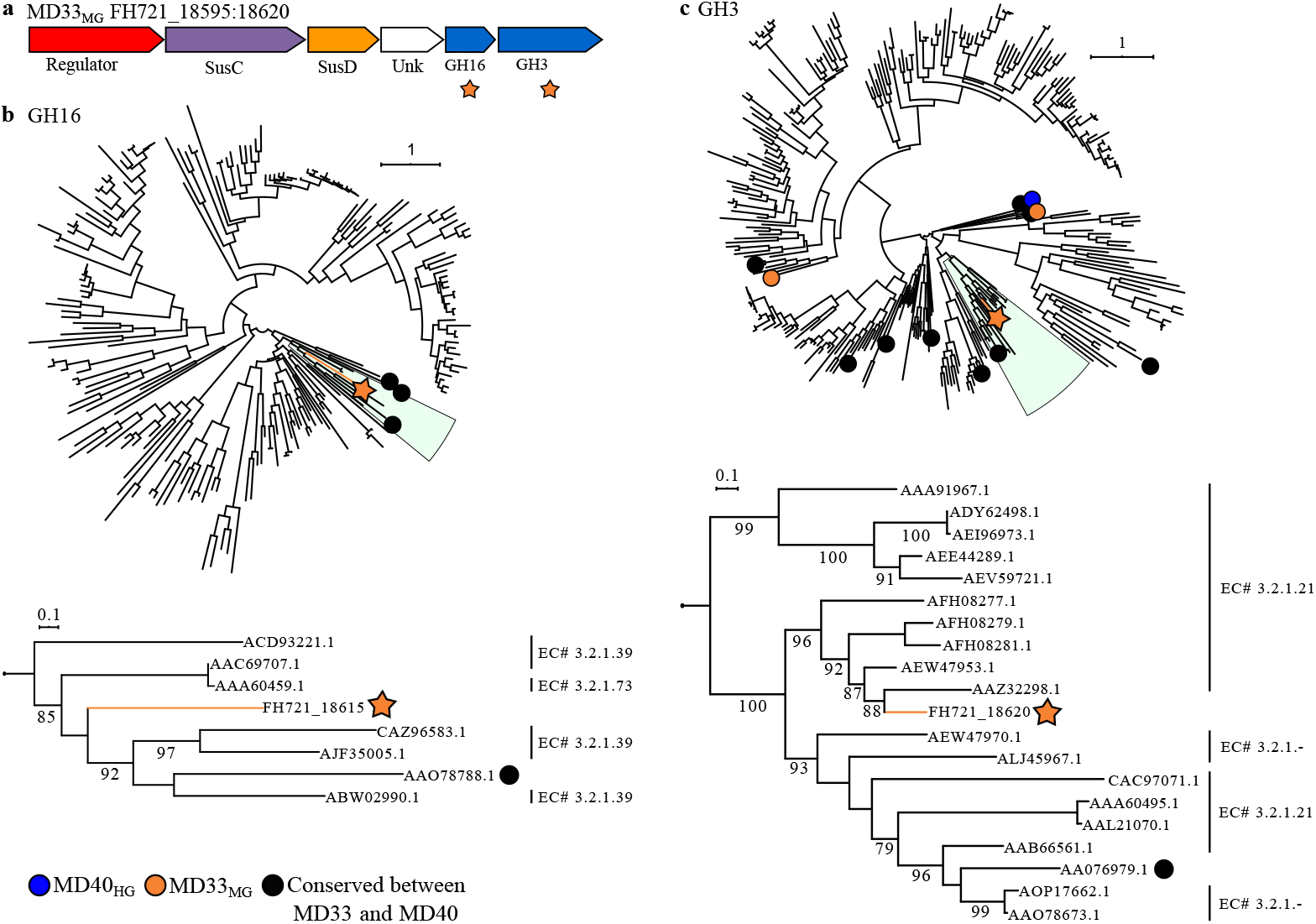
CAZome analysis of MD33_MG_ and MD40_HG_. Predicted PUL (FH721_18595-18620) found in the genome of MD33_MG_ encoding GH16 and GH3 enzymes, which are not found in the MD40_HG_ genome. Phylogenetic trees of GH16 (top) and GH3 (bottom) families in MD33_MG_ (orange) and MD40_HG_ (blue) genomes. (Left) Expanded clades with the unique GH16 and GH3 CAZymes are shown with known EC numbers.

## Materials & Methods

### Direct visualization of YM metabolism in rumen communities and cell identification by FISH

Rumen samples were collected from two cannulated cows fed a diet rich in barley grain. The rumen samples were filtered through cheesecloth under CO_2_ gas. Subsamples were taken, flash-frozen, and stored at −80°C until genomic extractions could be completed. The rest of the sample was transferred into an anaerobic chamber (atmosphere: 85% N_2_, 10% CO_2_, 5% H_2_, at 37°C) and filtered through a 100 μm pore size nylon net filter (Millipore, U.S.A). The filtered samples from each cow were then aliquoted into three tubes. One tube was immediately fixed with 1% formaldehyde (FA) for 1 h at room temperature as the 0 h control. The other tubes were incubated with 20 μL FLA-YM for a final concentration of 3.1 nM and fixed with FA after 1 d and 3 d. Immediately after fixation all samples were filtered through a 47 mm (0.2 μm pore size) polycarbonate filter (Millipore), using a 0.45 μm cellulose acetate support filter (Millipore) and a gentle vacuum of < 200 mbar. After drying, the filters were stored at −20°C.

Total cell counts were determined by staining with 4’,6-diamidino-2-phenylindole (DAPI) and visualizing on a Leica DMRX epifluorescence microscope (Leica, Germany). The number of FLA-YM stained cells were determined by enumerating cells which had a positive DAPI and FLA-YM signal (excitation at 405 nm and 488 nm wavelengths, respectively). For FISH, oligonucleotide probe CF968 targeting *Bacteroidetes* (5’-GGTAAGGTTCCTCGCGTA-3’) (Acinas et al., 2015) was used, which was covalently labeled with four ATTO594 fluorochromes by Biomers (Konstanz, Germany). FISH was performed with slight alterations to the protocol of Manz *et al* (1992) (Manz, Amann, Ludwig, Wagner, & Schleifer, 1992). The hybridization buffer contained 900 mM NaCl, 20 mM TRIS-HCl (pH 7.5), 0.02% sodium dodecyl sulfate, 10% dextran sulfate (wt/vol) and 1% (wt/vol) blocking reagent (Boehringer, Germany) with a formamide concentration of 55%.

Hybridizations were carried out at 35°C in a humidity chamber overnight, with a subsequent 15 min wash in a buffer containing 10 mM NaCl, 20 mM TRIS-HCl (pH 7.5), 5 nM EDTA (pH 8) and 0.01% sodium dodecyl sulfate at 37°C. After FISH, the abundance of *Bacteroidetes*, as well as, *Bacteroidetes* showing FLA-YM uptake was enumerated using a Leica DMRX epifluorescence microscope.

DNA from the frozen rumen samples was extracted using the Qiagen DNeasy PowerSoil Kit and samples were sent to McGill GenomeQuebec for Illumina MiSeq PE250 16S rDNA metagenomics sequencing. The 16S rDNA sequences were merged and quality trimmed using the BBTools(Bushnel) software and subsequently classified using the standard settings of the SILVAngs pipeline using the SSU rRNA seed of the SILVA database release 132 (Quast et al., 2013; Team, 2015). All analysis and plotting of the microbial diversity data were done using RStudio version 3.6.3 using the Vegan package (Oksanen et al., 2019; Team, 2019)

### Isolation of bovine-adapted mannan-degraders

Bovine rumen and fecal samples were collected for i*n vitro* batch culture experiments. Ruminal and fecal inoculants from cattle were enriched with one of the following substrates: Bio-Mos^®^ (1% W/V), corn distillers’ grains (1% W/V), or YM (1% W/V). Bacteria were isolated from the enriched batch cultures by streaking onto nutrient restricted media supplemented with 0.5% YM to select for YM-degraders (supplementary methods). In total, 50 YM-degrading bacterial isolates were characterized for their propensity to metabolize YM. Nine of these isolates were selected for detailed analysis in this study.

### Growth profiling of bovine isolates

Bovine isolates, wild type *Bt*VPI-5482, and a mutant *Bt*strain lacking MAN-PULs 1, 2, and 3 (ΔMAN-PUL1/2/3) (Cuskin, Lowe, et al., 2015a) were cultured overnight in tryptone-yeast-glucose (TYG) medium (supplementary methods). All incubations were performed in an anaerobic chamber at 37 °C. The overnight cultures (OD_600_ 1.0-1.4) were diluted to an OD_600_ of 0.05 in 2X *Bacteroides* minimal medium (MM), pH 7.2 (supplementary methods). Wells of a 96-well microtiter plates (Falcon) were filled with 100 μL of sterilized 1% (w/v) YM (Sigma, St. Loius, USA; M7504) or mannose along with 100 μL inoculant (n=4). Negative control wells consisted of 100 μL 2X MM combined with 100 μL 1% (w/v) of YM or mannose and were used to normalize growth curves. 100 μL of bacterial suspension was inoculated to get starting OD_600_~0.025. Plates were sealed with polyurethane Breathe-Easy gas-permeable membranes (Sigma; Z390059). Absorbance (600 nm) of each well was measured with a Biotek Eon microplate reader and recorded on Biotek Gen5 software every 10 min for 50 h. Mean (± standard deviation) of each condition (n = 4) was visualized using GraphPad Prism 6. Two replicates of each strain were also cultured on YM extracted from the cell wall of *S. pombe* (supplementary methods).

Post-growth cultures were harvested and centrifuged. Supernatants were taken and 6 μL was ran on a silica sheet in 2:1:1 (butanol:d_2_H2O:acetic acid) running buffer. The plate was dried at ambient temperature and stained with orcinol (diluted to 1% in a solution of 70:3 ethanol:sulfuric acid). Once the plate was dry, it was activated in an oven at 120°C and imaged using a gel doc XR image system (Bio-Rad).

### Genome sequencing, assembly, and annotation of *Bt*^Bov^ strains

The 16S rDNA of 50 bovine bacterial isolates was sequenced to determine taxonomic classification using the universal primers 27F and 1492R (supplementary methods). Based on growth profiles (OD_600_ > 0.4) and 16S rDNA sequences, nine isolates were chosen for whole genome sequencing using Illumina MiSeq PE150 bp. Genomes were assembled using SPAdes *de novo* assembly (Bankevich et al., 2012). The K-mer value in SPAdes was chosen from (21, 33, 55, 77 - defaults for 150 bp reads). Quality reporting of the assemblies was done using Quast (Gurevich, Saveliev, Vyahhi, & Tesler, 2013). SPAdes assembly N50s, largest contigs, and number of contigs are shown in S2 Table. Isolate contigs were blasted against the reference genome *Bt*VPI-5482 for MAN-PUL1/2/3, and the HMNG-PUL using NCBI BLAST (2.7.1) (Altschul, Gish, Miller, Myers, & Lipman, 1990). SPAdes contig assemblies were aligned with the JSpeciesWS reference *B. theta* genomes to calculate average nucleotide identity based on BLAST+ (ANIb) (GmbH, 2019; Richter et al., 2016).

### Production of *Bt*VPI-5482 MAN-PUL mutants

Flanking regions (~750bp) of the *susC/D*-like genes from each MAN-PUL were PCR amplified, stitched together, and ligated into pExchange-tdk (pEx-tdk). The plasmids were transformed into *E. coli* strain S17-1λpir, which were donor cells used to conjugate the plasmids into the *Bt*VPI-5482 ΔPUL75Δtdk recipient strain to delete the *sus*-like gene pairs (Jones et al., 2019). Mutants with the MAN-PUL1 Sus genes deleted (ΔMP1*susCD*) were then conjugated with *E. coli* cells containing a plasmid with the flanking regions for the MAN-PUL3 Sus genes to create a dual mutant (ΔMP1/3*susCD*). The dual mutant was then conjugated with *E. coli* cells that contained a plasmid with the MAN-PUL2 Sus flanks to produce a triple mutant(ΔMP1/2/3*susCD*). Plasmids and mutants were sequenced at each step of this process.

The three knock-out strains, along with *Bt*VPI-5482 wild-type, were grown on 0.5% YM-MM as described above. In addition, the strains were incubated with FLA-YM, and sampled at 0hr, 1hr, 1d, and 3d. These samples were fixed and stored at 4°C until analyzed by flow cytometry and epifluorescence microscopy (see below).

### PUL delineation and comparative CAZomics

Isolate contigs were processed through EMBOSS GetORF (Rice, Longden, & Bleasby, 2000) to determine open reading frames; these data were run through the dbCAN (Yanbin Yin et al., 2012) HMMscan to identify CAZyme sequences. CAZyme sequences were then analyzed by SACCHARIS (Jones et al., 2018) (<u>S</u>equence <u>A</u>nalysis and <u>C</u>lustering of <u>C</u>arbo<u>H</u>ydrate <u>A</u>ctive enzymes for <u>R</u>apid <u>I</u>nformed prediction of <u>S</u>pecificity). User CAZyme sequences were trimmed to their catalytic domain with dbCAN(Y. Yin et al., 2012), aligned with MUSCLE(Edgar, 2004), and fitted to a phylogenetic tree using ProtTest3 (Darriba, Taboada, Doallo, & Posada, 2011) to find the appropriate amino acid replacement model. RAxML (Stamatakis, 2014) or FastTree (Price, Dehal, & Arkin, 2010) was used to generate the final tree. GHs from families 38, 76, 92, 99, and 125 identified in the genomes of the MD isolates were analyzed by SACCHARIS. Phylogenetic trees were developed using FastTree, and Newick file outputs were viewed using FigTree (Rambaut, 2009) and plotted using ITOL.

### RNA-seq: assembly, quantitation, and comparative analysis

RNA from *Bt*VPI-5482, MD33_MG_, and MD40_HG_ grown in 1% mannose or YM (see supplementary methods) was extracted and purified using a GeneJET RNA Purification kit (Thermo Scientific) within 1 week of storage. RNA was sent to Génome Québec for Illumina HiSeq 4000 PE100bp sequencing. Using Geneious v11.1.2(Kearse et al., 2012), each set of reads was mapped to their previously assembled genomic sequence, or in the case of *Bt*VPI-5482, to the genomic sequence from the NCBI database (NC_004663). Expression levels were calculated as transcript expression (transcript per kb per million; TPM) for each growth treatment. Ambiguously mapped reads were counted as partial matches. The Geneious DESeq2(Love, Huber, & Anders, 2014) plugin was used to compare the expression levels between the two treatments, producing log_2_ expression ratios and p-values.

Generalized linear mixed models in SAS PROC GLIMMIX (SAS 9.4, SAS Institute, Cary, NC) were used to estimate statistically significant (*p* < 0.05) differences of TPM means (least squares-means) for the MAN-PUL1/2/3 *susC*-like genes of each bacterial strain. Based on the Bayesian information criterion (BIC) of the generalized linear mixed models, the response was modeled using the log-normal distribution. The expression of gene transcripts was the dependent variable in models with two independent fixed factors: bacterial strain (*i.e*. *Bt*VPI-5482, MD33_MG_ or MD40_HG_) and media treatment (*i.e*. YM or mannose). Mixed models of variance heterogeneity were selected based on the BIC. For the studied transcripts, the variance of expression was heterogeneous for the experimental treatments, bacteria, or their interaction. The statistical significance of the interaction between the TPM values of MANPUL genes for each bacterial strain and the media treatment was determined using an *F*-test. Bonferroni’s method was used for multiple comparisons (S3 Table).

### Production of SusD-like protein C-myc fusion *B. theta* strain

The C-myc epitope (EQKLISEEDL) was fused to the C-terminal domain of the MAN-PUL2 SusD-like protein (BT_3789) with a linker sequence (STSTST) between the SusD-like nucleotide sequence and the C-myc sequence *Bt*VPI-5482 Δtdk Δpul75 (control) and *Bt*VPI-5482 Δtdk Δpul75 SusD-like C-myc fusion mutant were inoculated in TYG and cultured as described above. The cells were centrifuged and resuspended in 1 mL 2X MM. 100 μL of the resuspension was inoculated into 0.5% YM-MM and incubated for 4 h at 37°C. The cells were then centrifuged and washed three times in phosphate-buffered saline (PBS) pH 7.4 (PBS; 137 mM NaCl, 2.7 mM KCl, 10 mM Na_2_HPO_4_), before resuspension in 2 mL 2X MM. 225 μL of the resuspended cells were added to 1.5 mL 0.2% FLA-YM or YM-MM and incubated for 3 h at 37°C. 100 μL of each culture was collected and fixed in 1% formaldehyde for 1 h at room temperature. The samples were then incubated with 1:2500 rabbit IgG anti-C-myc polyclonal antibody for 1 h at room temperature. The samples were then washed four times in PBS and resuspended in 1:2500 goat anti-rabbit DyLight 650nm secondary antibody for 1h at room temperature. The samples were then washed and stored in PBS until further analysis.

### Sequence comparison and modelling of SusC/D/E-like proteins

MUSCLE was used to align MAN-PUL2 and 3 SusC-like, SusD-like (SGBPA), and SusE-like (SGBPB) amino acid sequences of each isolate and calculate percent identity (S4 Table). The 16S rDNA gene and MAN-PUL2 SusC-like and SusD-like amino acid phylogenetic trees were generated using the Maximum Likelihood method and Tamura-Nei model(Tamura & Nei, 1993). Evolutionary analyses were performed by MEGA X(Kumar, Stecher, Li, Knyaz, & Tamura, 2018). Trees with the highest log likelihood are shown in Fig. 4b,c.

### Generation of FLA-YM conjugates

A previously defined protocol (Arnosti, 2003; Reintjes et al., 2017) was used to generate fluorescently labelled YM (FLA-YM), with slight variations (supplementary methods).

### Visualization of FLA-YM uptake by strains of *Bt*^Bov^

Wild type *Bt*VPI-5482, *Bt*ΔMAN-PUL1/2/3, and rumen isolates MD33_MG_, and MD40_HG_ were inoculated in TYG and grown as described above. Cells were harvested at OD_600_ ~1.0 and centrifuged (4,700 x *g*) for 5 min, the supernatant was removed, and pellets resuspended in 2 mL 2X MM for the first two washes. After the third centrifugation, pellets were resuspended in 2 mL MM with 0.5% YM *fBt*VPI-5482, MD33_MG_, and MD40_HG_) or 0.5% glucose + YM (*Bt*ΔMAN-PUL1/2/3) as the sole carbon source (not conjugated to FLA). After ~18 h incubation, cultures were centrifuged and washed three times in PBS, with the final resuspension in 2 mL 2X MM. 300 μL of the resuspended pellet was aliquoted into 0.2% unlabeled YM or FLA-YM. 20 μL of the 2X MM resuspension was used as the 0 h time point, as the cells were not exposed to FLA-YM. 40 μL aliquots of each condition were taken at time points: 5 min, 1 h, and 24 h. The cells were centrifuged (10 min; 2,300 x *g*) and the pellet was fixed in 1% formaldehyde (FA; Sigma; F8775) in PBS, at 4°C for 18 - 24 h. The fixed cells were centrifuged (10 min; 2,300 x *g*) and washed in 1 X PBS. The samples were centrifuged and stored at 4°C in the dark until visualized by SR-SIM (supplementary methods).

### Quantification of the rate of FLA-YM uptake by *Bt*^Bov^ isolates

*Bt*VPI-5482, MD33_MG_, and MD40_HG_ were grown in TYG and prepared as described above. After 24 h of incubation, cultures were placed into 2 mL 0.5% YM. Cells were harvested in exponential phase (OD_600_ 0.6-1.0), centrifuged (10 min; 2,300 x *g*), and resuspended in 2 mL 2X MM. 300 μL of this suspension was added to 1 mL 2 X MM. Then 20 μL of each culture was aliquoted into 1 mL 1% FA and used as the T0 time point. Into the remaining 280 μL, 0.2% FLA-YM, 150 ng/mL fluoresceinamine (FLA) or YM was added and subsamples of 40 μL were taken at 5, 10, 15, 20, 30 and 60 min, 2, 4, 8 and 24 h. The subsamples were centrifuged (10 min, 2,300 x *g*) and the cell pellets were fixed in 1% FA in 1X PBS, at 4°C for 18 – 24 h. The fixed cells were centrifuged (10 min; 2,300 x *g*) and resuspended in 1 ml 1X PBS and stored at 4°C in the dark.

Cell fluorescence due to FLA or FLA-YM uptake was quantified in all samples using an Accuri C6 flow cytometer (BD Accuri Cytometers). The 8-peak and 6-peak validation bead suspensions (Spherotech, IL, USA) were used as internal references. All samples were measured under laser excitation at 488 nm from a blue-green laser and the green fluorescence was collected in the FL1 channel (530 ± 30 nm). Using the medium as a background an electric threshold of 17,000 FSC-H was set to reduce the background noise. All measurements were done at a slow flow rate and a total of 10,000 (FLA-YM) or 5,000 (YM and FLA) events per sample were acquired. Bacteria were detected from the signature plot of SSC-H vs green fluorescence (FL1-H). The FCM output was analyzed using FlowJo v10-4-2 (Tree Star, USA). The FCM files were imported into FlowJo and both the total population (all events) and main population (automated gating through event density) were determined. For each population (total and main) sample statistics (counts, mean fluorescence and the standard deviation) were determined from the raw FL1-H data. The results were exported and analyzed using Welch’s *t*-tests in R studio using the packages Vegan and Rioja (Juggins, 2016; Oksanen et al., 2013; Team, 2015) to determine statistical difference between the control (YM and FLA) and FLA-YM incubation within each strain, and between the FLA-YM incubation of each strain.

### Quantification of mannose in minimal medium using GC-MS

Cell culture medium after incubation in 1 % YM-MM was collected after 24 hrs, centrifuged (4,700 x *g* for 15 mins), and the supernatant was passed through a syringe filter (0.2 μm cellulose acetate membrane, VWR). The filtrate was kept frozen for 48 hrs at −20°C and then thawed and centrifuged (3000 x *g*, 30 min) at room temperature. Concentration of mannose in the resulting supernatant was tested based on our previous report (MacMillan et al., 2019), with some modifications to cope with the relatively large amount of starting carbohydrate material and the presence of minimum medium. One milliliter of the supernatant was evaporated to dryness under a gentle flow of nitrogen. The residue was suspended and magnetically stirred in 3.5 mL of 6 M TFA at 100°C for 6 hrs with headspace filled with nitrogen, followed by addition of internal standard myo-inositol (0.4 mg dissolved in 0.5 mL of water) and evaporation to dryness. Monosaccharides were reduced by magnetic stirring overnight in 10 mg of NaBD4 (99% D, Alfa Aesar) dissolved in 2 mL of 1 M ammonium oxide solution, followed by quenching excess reductant with acetic acid and evaporating to dryness. Boric acid was removed by evaporation to dryness five times in 3 mL of 10% (v/v) acetic acid in methanol followed by five times in 3 mL of absolute methanol. The residue was suspended in 4 mL of acetic anhydride, followed by magnetic stirring at 100°C for 2 hrs with headspace filled with nitrogen, cooling to room temperature, and evaporation to dryness. The derivatives were purified by partitioning with water and dichloromethane, recovered by collecting and evaporating to dryness the organic phase after three changes of water, and re-dissolved and diluted in ethyl acetate for analysis on an Agilent 7890A-5977B GC-MS system (Agilent Technologies, Inc., CA). Sample solution (1 μL) was splitless-injected to the system, and optimal analyte separation was achieved on a medium polarity SP2380 column (30 m × 0.25 mm × 0.20 μm, Sigma-Aldrich) with a constant helium flow of 0.8 mL/min and with oven temperature programmed to start at 55°C (hold 1 min) followed by increasing at 30°C/min to 120°C then at 12°C/min to 255°C (hold 20 min). Two separate experiments were conducted for each sample. Mannose concentration was calculated based on calibration curve established from a series of mannose standard solution containing internal standard.

### Measurement of YM Hydrolysis

Samples from *Bt*VPI-5482, MD33_MG_, and MD40_HG_ cultures in 0.2% FLA-YM (as above) and a no cell negative control were filtered through 0.2 μm cellulose acetate membrane syringe filter (VWR). The filtrates were flash frozen and stored at −80°C until analysis. Samples were analyzed as described in Arnosti (Arnosti, 2003); in brief, samples were injected onto two columns of Sephadex G50 and G75 gel linked in series, with the column effluent passing through a Hitachi fluorescence detector set to excitation and emission wavelengths of 490 and 530 nm, respectively. The columns were standardized using FITC dextran standards (>50 kDa, 10 kDa, 4 kDa, monosaccharide, and free fluorophore), so the fraction of polysaccharide eluting in each molecular weight class at each time point could be calculated.

## Supporting information

Supplementary Figures Klassen and Reintjes 2020

Supplementary Material Klassen and Reintjes 2020

## Acknowledgements

This work was supported by funding from the Beef and Cattle Research Council awarded to DWA (Grant: FDE.13.15 & FDE.14.17). JHH, GR, and RA were supported by the Max Planck Society. RA and JHH acknowledge support by the Deutsche Forschungsgemeinschaft (DFG) in the framework of the research unit FOR2406 ‘Proteogenomics of Marine Polysaccharide Utilization (POMPU)’ by grants AM 73/9-1 and HE 7217/1-1. GR has received funding from the European Union’s Horizon 2020 research and innovation program under the Marie Sklodowska-Curie grant agreement No. 840804. CAr was supported by NSF OCE-1736772, DE-SC0013887, and DE-SC0019012. We wish to thank Sherif Ghobrial and Chad Lloyd for assistance with the FLA-YM hydrolysis measurements, and Elizabeth Lowe, Newcastle University for the kind gift of the MAN-PUL1/2/3 KO mutant.

## Author Contributions

LK assisted with rumen extractions and performed rumen incubations with FLA-YM, selective bacterial growth profiling and TLC analysis, RNA sequencing, production of FLA-YM, FGC-bacterial incubations, construction of the SusD-C-myc mutant and evaluation of production, extraction of *S. pombe* YM, prepared figures, and wrote manuscript. GR conducted epifluorescence microscopy and SR-SIM analysis, FISH analysis, cell enumeration of rumen samples, flow cytometry sorting and analysis, statistical analysis and assisted with figure and manuscript writing and preparation. JPT performed comparative genome and protein sequence analysis, and assisted with figure preparation. DRJ assisted with comparative genome and protein analysis. JHH helped conceive of the study, interpreted data, and assisted in the preparation of the manuscript. ADS performed isolations of rumen bacteria and 16S sequencing. TDS conducted statistical analysis of RNAseq data. CAr assisted with generation of FLA-PS, analysis FLA-PS products, and preparation of the manuscript. LJ performed rumen collections and assisted with bacterial isolations. TWA assisted with rumen extractions and preparation of the manuscript. CAm assisted with RNAseq analysis and figure generation. DT assisted with CAZyme fingerprinting and maintenance of the SACCHARIS pipeline. RA helped conceive of the study, assisted with SR-SIM microscopy and preparation of the manuscript. TAM assisted with animal study and rumen extractions, and preparation of the manuscript. DPY assisted with data analysis and preparation of the manuscript. DWA helped conceive of the study, secured funding, designed the study, performed data analysis, and assisted with figure and manuscript preparation.

