## Supplementary Figures Klassen and Reintjes 2020 for "Quantifying fluorescent glycan uptake to elucidate strain-level variability in foraging behaviors of rumen bacteria"

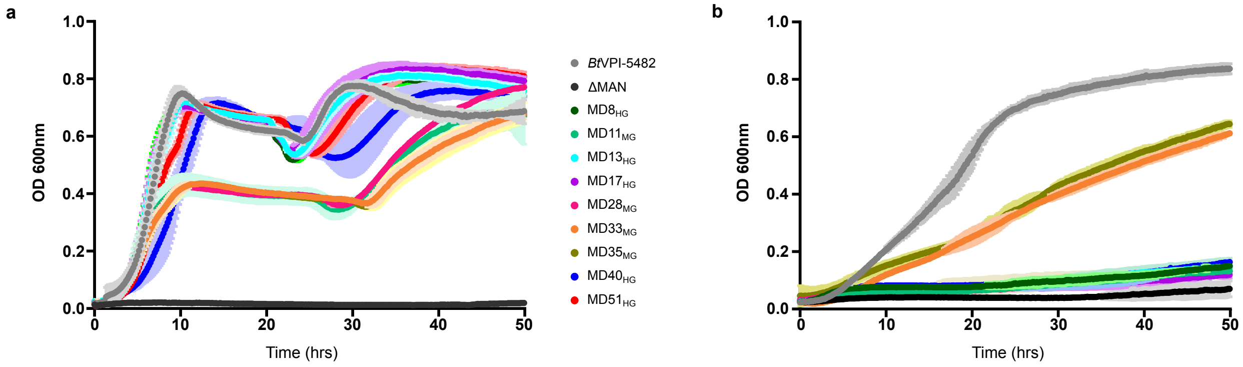


Supplemental Figure 1, Klassen and Reintjes et al., 2020

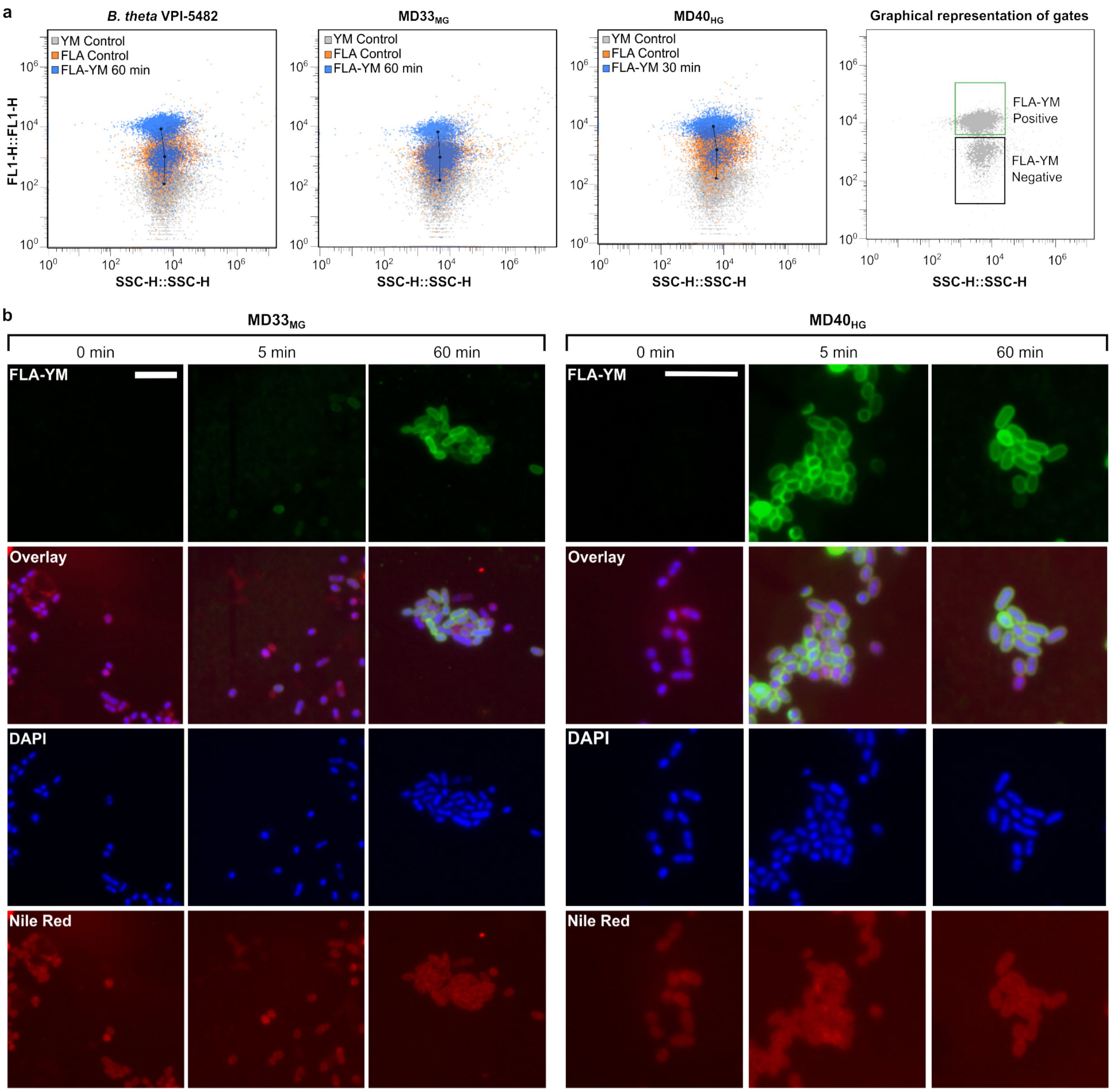


Supplemental Figure 2, Klassen and Reintjes et al., 2020

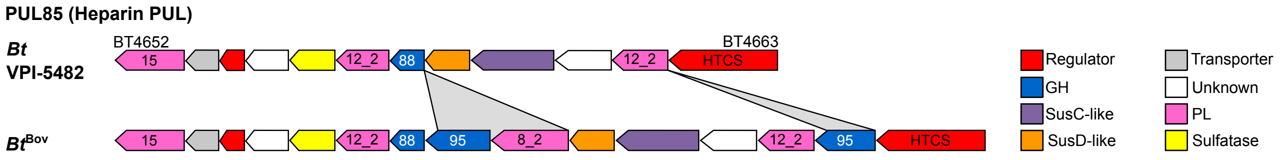


Supplemental Figure 3, Klassen and Reintjes et al., 2020

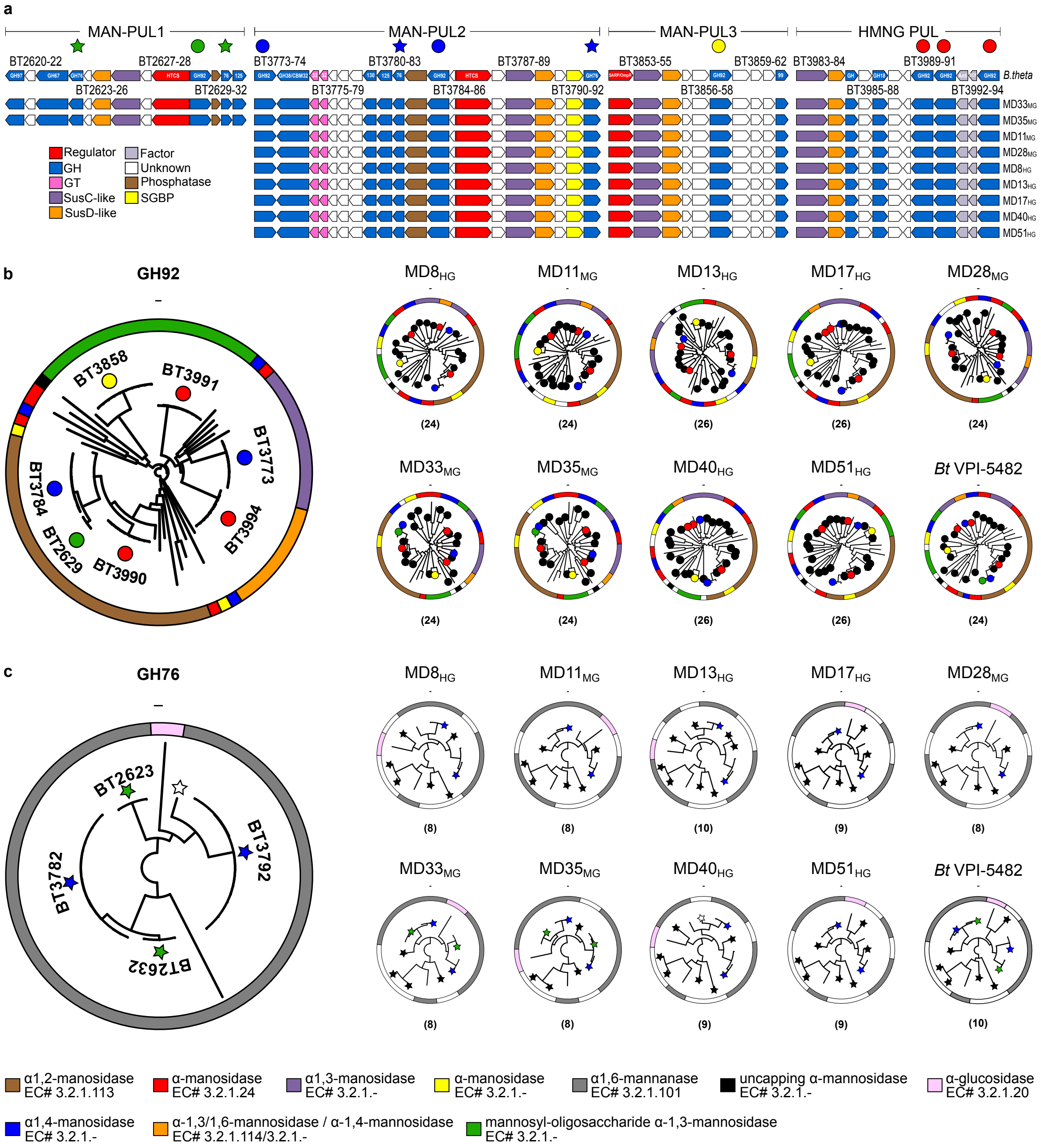


Supplemental Figure 4, Klassen and Reintjes et al., 2020

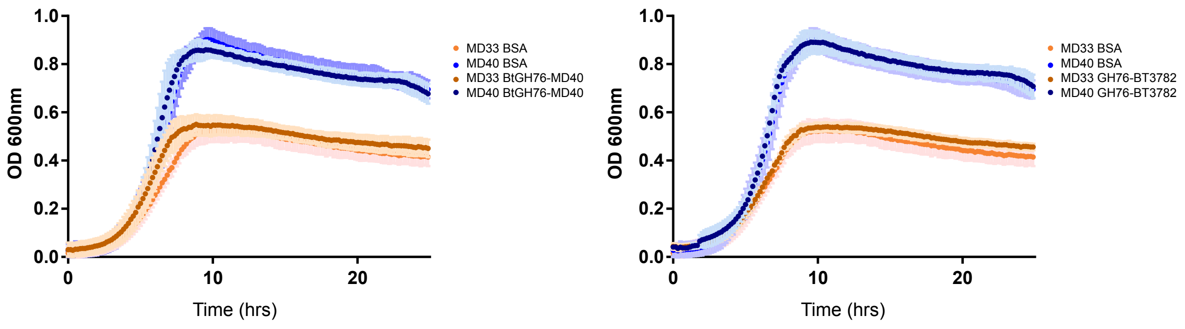


Supplemental Figure 5, Klassen and Reintjes et al., 2020

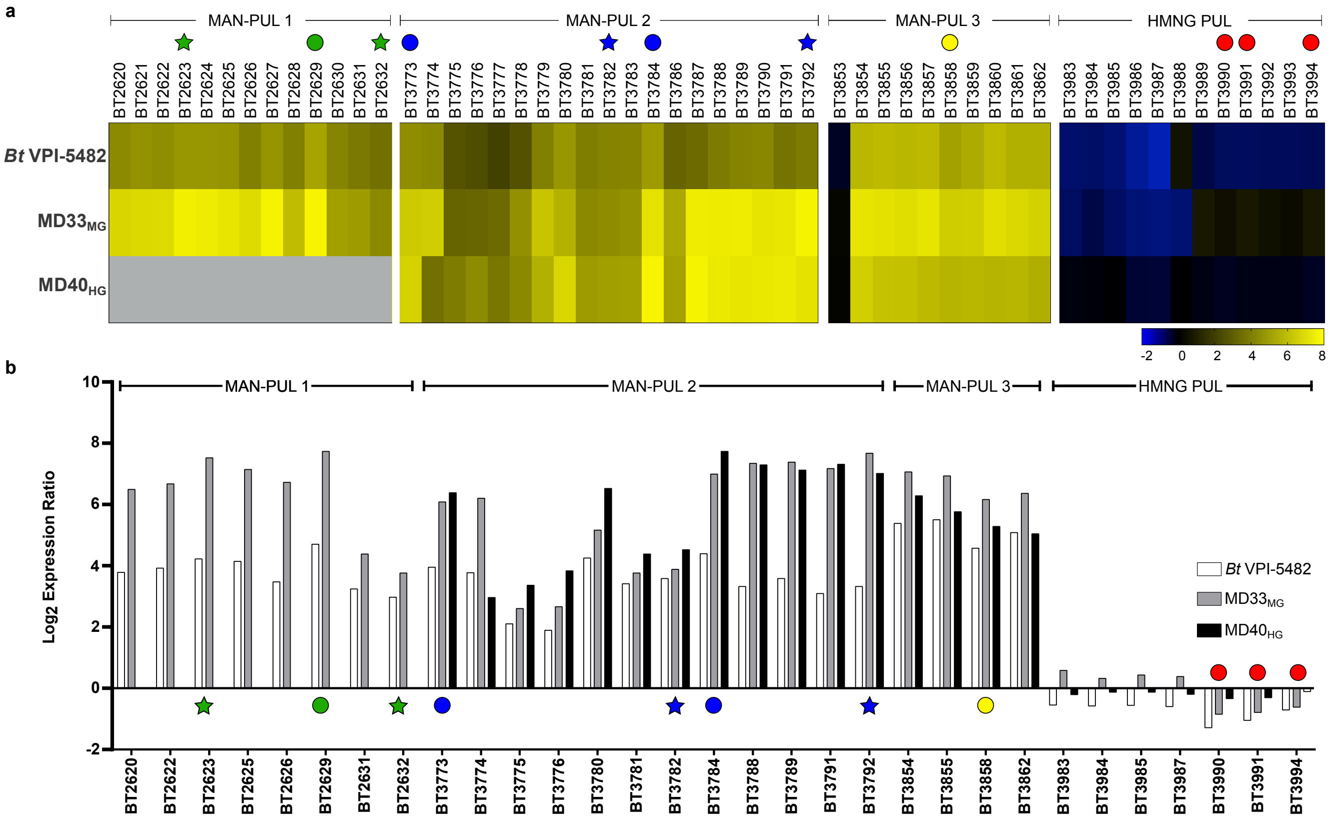


Supplemental Figure 6, Klassen and Reintjes et al., 2020

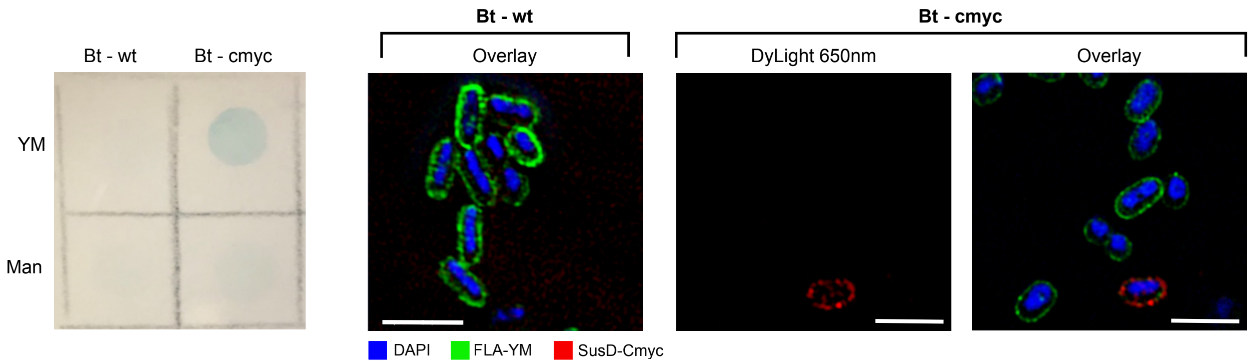


Supplemental Figure 7, Klassen and Reintjes et al., 2020

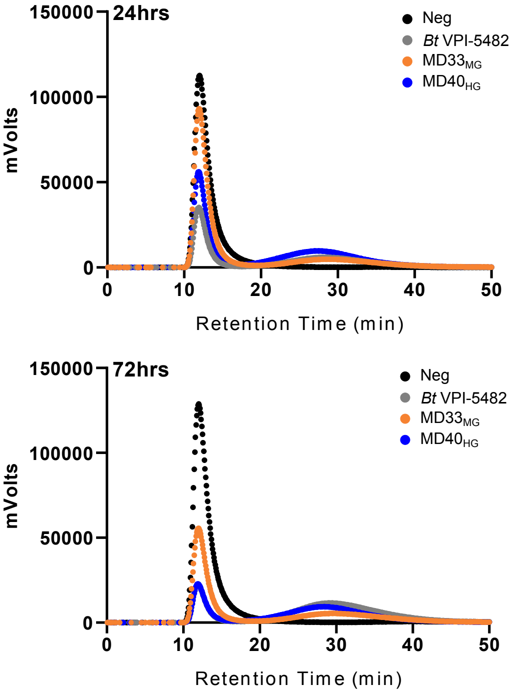


Supplemental Figure 8, Klassen and Reintjes et al., 2020
