## Supplementary Material Klassen and Reintjes 2020 for "Quantifying fluorescent glycan uptake to elucidate strain-level variability in foraging behaviors of rumen bacteria"

### Supplementary Discussion

#### Catabolite repression in *B. theta* when grown on YM.

In addition to MAN-PUL activation, regions of gene repression were observed in each strain (Fig. 5a). *Bt*VPI-5482 down-regulated ($log_2$ fold-change < -3.5) many genes when grown on YM, including PUL22, which is predicted to target dietary fructans (*BT_1758-1763*)(Martens, Chiang, & Gordon, 2008; Sonnenburg et al., 2010). This response was observed in $MD33_{MG}$ and $MD40_{HG}$ for a homologous cluster of fructan-utilizing genes ($log_2$ fold-change < -3.5), suggesting there may be a YM-catabolite repression effect with these two polysaccharides, as it was not observed when the cultures were grown on mannose. In MAN-PUL2, *BT_3775-BT_3776*, which encodes a biosynthetic gene cluster, displayed the lowest expression levels of all the MAN-PUL genes. This is consistent with what was reported previously (Martens et al., 2011). Overall, the $MD33_{MG}$ transcriptome had more plasticity in the genes that were either induced or repressed in comparison other two strains (Fig. 5a).

#### Expression levels of MAN-PULs

Within each MAN-PUL, the *gh*, *susC*-like, and *susD*-like gene clusters consistently displayed the highest $log_2$ expression values (Fig. 5b, Supplementary Fig. 6). The BT_3788 (*susC*-like, 5.4) and BT_3789 (*susD*-like, 5.5) genes from MAN-PUL3 in *Bt*VPI-5482, BT_2629 (*gh92*, 7.7) and BT_3792 (*gh76*, 7.7) genes in $MD33_{MG}$, and BT_3784 (*gh92*, 7.7) gene in $MD40_{HG}$ displayed the highest expression of all genes studied (Supplementary Fig. 6a).

#### Host adaptation of $Bt^{BOV}$ strains

We observed the presence of signature CAZymes that suggested these strains have adapted to colonize the bovine gut. For example, in every $Bt^{Bov}$ strain, two GH95s and two polysaccharide lyases from Family 8 are inserted into the host-glycan utilization locus PUL85, which differs from the human-associated *Bt*VPI-5482 and 7330 strains (Supplementary Fig. 3). If such host-bacterium interactions are factors required for host colonization, these properties may be exploited for developing bovine-adapted probiotics.

### Supplementary Methods

#### Isolation of bovine-adapted mannan-degraders

Three biological replicates for each sample were incubated anaerobically at 39˚C on a rotary shaker for 24 h. Serial dilutions of the batch cultures ($10^{0}$-$10^{-6}$) were similarly streaked on YM plates. Plates were incubated anaerobically (atmosphere: 85% $N_2$, 10% $CO_2$, 5% $H_2$) at 37˚C for up to 96 h. After 24-96 h of incubation, single colonies were selected and inoculated in nutrient rich Brain Heart Infusion (BHI) medium: 3.7% Bacto™ Brain Heart Infusion (BD; 237500), 8.3 mM L-cysteine, 10mL/L hemin (0.77 mM hemin in 0.01M NaOH), and 0.2% $NaHCO_3$. Overnight cultures were centrifuged and resuspended in 0.8 mL glycerol (50%) and stored at -80˚C.

#### Growth profiling of bovine isolates

TYG Ingredients: 1% Bacto™ Tryptone (BD; 211705), 0.5% Yeast Extract Bacteriological (VWR; J850), 4.1 mM L-cysteine, 0.2% glucose, 0.1 M $KPO_4$ pH 7.2, 2.2 µM vitamin $K_3$, 40 µL/mL TYG Salts (2 mM $MgSO_4 \cdot 7H_2O$, 119 mM $NaHCO_3$, and 34.2 mM NaCl), 28.8 µM $CaCl_2$, 1.4 µM $FeSO_4$, 4.4 µM resazurin, 1 µL/mL (v/v) histidine/hematin (1.9 mM hematin, and 200 mM L-histidine, 1000X stock solution).

*Bacteroides* MM Ingredients: : 200 mL/L (v/v) 10X Bacteroides salts solution pH 7.2 (999 mM $KH_2PO_4$, 150 mM NaCl, 85 mM $(NH_4)_2SO_4$), 20 mL/L (v/v) Balch's Vitamins pH 7.0 (36.5 µM ρ-aminobenzoic acid, 4.5 µM folic acid, 8.2 µM biotin, 40.6 µM nicotinic acid, 10.5 µM calcium pantothenate, 13.3 μM riboflavin, 14.8 μM thiamine HCl, 48.6 μM vitamin $B_6$, 73.8 nM vitamin $B_{12}$, and 24.2 mM thioctic acid), 20 mL/L (v/v) Amino Acid Solution (5 mg/mL amino acids: alanine, arginine, asparagine, aspartic acid, cysteine, glutamic acid, glutamine, glycine, histidine, isoleucine, leucine, lysine, methionine, phenylalanine, proline, serine, threonine, tryptophan, tyrosine, and valine), 20 mL/L (v/v) Purine/Pyrimidine Solution pH 7.0 (1 mg/mL adenine, guanine, thymine, cytosine, and uracil), 20 mL/L (v/v) Trace Mineral Solution pH 7.0 (1.7 mM ethylenediaminetetraacetic acid, 12.2 mM $MgSO_4 \cdot 7H_2O$, 3 mM $MnSO_4 \cdot H_2O$, 17.1 mM NaCl, 359.7 μM $FeSO_4 \cdot 7H_2O$, 901.1 μM $CaCl_2$, 347.7 μM $ZnSO_4 \cdot 7H_2O$, 40.1 μM $CuSO_4 \cdot 5H_2O$, 161.7 μM $H_3BO_3$, 41.3 μM $Na_2MoO_4 \cdot 2H_2O$, and 84.1 μM $NiCl_2 \cdot 6H_2O$), 4.4 μM vitamin $K_3$, 2.9 μM $FeSO_4 \cdot 7H_2O$, 14.4 μM $CaCl_2$, 2 mM $MgCl_2 \cdot 6H_2O$, 7.4 pM vitamin $B_{12}$, 16.5 mM L-cysteine, and 2 μL/mL (v/v) histidine/hematin.

**Extraction of YM from the cell wall of *S. pombe***

*S. pombe* was grown in 1 L Yeast Peptone Tryptone (YPD) medium at 30˚C with shaking at 150 rpm for 24 hours. Cells were harvested by centrifugation at 6,000 x g for 10 min. Pellets were resuspended in 10 mL 20 mM Citrate Buffer, pH 7.0 and autoclaved at 125˚C for 90 min. Autoclaved cells were centrifuged at 10,000 x g for 10 min. Supernatant was stored at 4˚C. Pellets were again resuspended in Citrate Buffer, autoclaved, and centrifuged. Supernatants were pooled and mixed with an equal volume of Fehling's Reagent. The solution was incubated at 40˚C, 120 rpm for 2 h. White precipitate formed in the solution and was harvested by centrifugation at 5,000 x g for 10 min. Pellets were dissolved in 2 mL 3 M HCl (per pellet from 1 L of original culture). The solution was slowly transferred to 100 mL of 8:1 Methanol:Acetic Acid with gentle stirring and incubated for 15 min. White precipitate formed in the solution and was collected by centrifugation at 3,200 x g. Supernatant was discarded and the pellet was dissolved in 100 mL 8:1

Methanol:Acetic Acid, vortexed briefly, and centrifuged at 1,500 x g for 5 min. This wash step was performed two more times and then repeated an additional three times with 100% Methanol. After the final wash, the pellets were left to dry. The following day, the pellets were dissolved in Mq $H_2O$ and dialyzed (500-1,000 MWCO) in Mq $H_2O$ for 20 h. The dialyzed solution was then freeze dried and used for growth analysis.

**Genome and 16S rDNA gene sequencing, assembly, and annotation of *Bt*$^{\mathbf{Bov}}$ strains**

The 16S rDNA was PCR amplified using the following two primers:

Universal primer 27F: 5’-AGR GTT TGA TCM TGG CTC AG-3’

Universal primer 1492R: 5’-GGT TAC CTT GTT ACG ACT T-3’

**RNA-seq: assembly, quantitation, and comparative analysis**

*Bt*VPI-5482, $MD33_{MG}$, and $MD40_{HG}$ were each inoculated into three tubes containing 5 mL of TYG media. Overnight cultures of *Bt*VPI-5482, $MD33_{MG}$, and $MD40_{HG}$ ($OD_{600}$ 1.0-1.4) were diluted with 2X MM to an $OD_{600}$ 0.05. 100 µL of 1% YM or mannose was added to six wells of diluted culture aliquoted from the same overnight stock, to a final volume of 200 µL. This step was repeated three times for a total of three replicates (each divided among six wells) for each bacterial strain and treatment. Cells were harvested during the exponential phase of the first growth phase ($OD_{600}$ 0.4-0.8). Six wells were pooled and added to an equal volume of RNAprotect (Qiagen). The mixture was vortexed for 5 sec and incubated for 5 min at room temperature. After incubation, the protected cells were centrifuged (10 min; 2,800 x *g*). The supernatant was decanted and pellets were stored at -80°C until further processing.

**Production of SusD-like protein C-myc fusion *B. theta* strain**

The 5' and 3' regions flanking the *BT_3789* stop codon were amplified by PCR. Primers that contained the linker+C-myc nucleotide sequences were used to amplify the PCR products. These amplicons were then stitched together and ligated into a pEXchange plasmid. The plasmid was transformed into *E. coli* and conjugated into *Bt*VPI-5482 Δtdk Δpul75 recipient strain. Insertion of the C-myc tag was confirmed by sanger sequencing.

**Generation of FLA-YM conjugates**

To chemically activate the polysaccharide, 350 µL 0.81 M cyanogen bromide (CNBr; 97%; Sigma C91492) was added to 2 mL 2% YM. For ~5 min, pH was monitored and maintained above 9.5 with additions of 0.25 M NaOH. Activated YM was separated from CNBr using Sephadex® G-50 gel filtration medium in a column coupled to a Bio-Rad BioLogic LP Multistatic peristaltic pump (flow rate: 1 mL $min^{-1}$). The mobile phase consisted of 0.2 M sodium tetraborate decahydrate pH 8.0 (≥99.5%, Sigma S9640). Activated YM was eluted into a vial containing 2.0 mg fluoresceinamine Isomer II (FLA; ~95%; Sigma O7985) wrapped in aluminum foil and incubated for ~24 h at room temperature. To remove excess FLA and purify labelled YM, the reaction mixture was loaded onto Sartorius Vivaspin 15R columns (5,000 MWCO; VS15RH11) and centrifuged (210 x *g*). Columns were repeatedly topped up with distilled $H_2O$ and centrifuged until a clear filtrate was observed. The purified FLA-YM was lyophilized, covered in aluminum foil and stably stored (~4°C) until further use.

**Visualization of FLA-YM uptake by strains of *Bt*^Bov^**

For microscopy, samples were resuspended in 1 ml 1 X PBS and, subsequently, 25 µl was heat fixed at 40°C onto a poly-L-lysine coated glass cover slip (12 mm, #1.5H) (Thorlabs GmbH,

Germany). After heat fixation the samples were washed in MQ to remove additional salts and dried at 35°C. The samples were then counter stained with 4', 6-diamidino-2-phenylindole (DAPI) (1 ng µl$^{-1}$ WS) and Nile red (2 ng µl$^{-1}$ WS) for 10 and 25 min, respectively. After each stain, the cells were washed in MQ and subsequently left to dry at RT. Finally, they were mounted onto glass slides using a 4:1 Citifluor (Electron Microscopy Sciences, USA) / VectaShield (Vector Laboratories, Germany) mounting solution.

For epifluorescence microscopy, the samples were visualised using a Zeiss Axioskop 2 motplus fluorescence microscopy with a 100x oil objective. The cells were imaged using the Axiovision software (Zeiss, Germany) and a constant exposure time of 200 ms was applied for the FLA-YM channel (Alexa Fluor 488 filter cube) to enable a comparison of the FLA-YM signal. The subcellular localisation of FLA-YM within individual cells was achieved using a super-resolution structured illumination microscopy (SR-SIM). A Zeiss ELYRA PS.1 microscope with 561, 488, 405 nm lasers and BP 573-613, BP 502-538, and BP 420-480 + LP 750 optical filters was used. Z-stack images were taken with a Plan-Apochromat x63/1.4 oil objective and processed with the ZEN2011 software (Carl Zeiss, Germany).

**S1 Table: Alignment of 16S rDNA gene sequences from mannan-degrading *B. theta* strains isolated from the rumen blasted against the NCBI database(Altschul, Gish, Miller, Myers, & Lipman, 1990).**

| Strain | Top Blast Hit(s) | Query cover (%) | Identity (%) |
|---|---|---|---|
| **$MD11_{MG}$** | *B. theta* VPI-5482 and strain 7330 | 100 | 99.7 |
| **$MD28_{MG}$** | *B. theta* strain 7330 | 100 | 99.7 |
| **$MD33_{MG}$** | *B. theta* VPI-5482 | 100 | 99.8 |
| **$MD35_{MG}$** | *B. theta* VPI-5482 and strain 7330 | 100 | 99.7 |
| **$MD8_{HG}$** | *B. theta* VPI-5482 and strain 7330 | 100 | 99.7 |
| **$MD13_{HG}$** | *B. theta* VPI-5482 and strain 7330 | 100 | 99.7 |
| **$MD17_{HG}$** | *B. theta* strain 7330 | 100 | 99.7 |
| **$MD40_{HG}$** | *B. theta* strain DMF | 100 | 100 |
| **$MD51_{HG}$** | *B. theta* strain 7330 | 100 | 100 |

**S2 Table: NCBI accession number and assembly parameters of bovine-associated bacterial isolates.** SPAdes *de novo* genome assembly output of nine $Bt^{Bov}$ isolates sequenced by Illumina MiSeq PE150bp and ANIb output of $Bt^{Bov}$ assembled contigs blasted to multiple *B. theta* strains from the JSpeciesWS genome reference database.

| ACCESSION NUMBER | ISOLATE | SOURCE | SPADES CONTIGS | SPADES LARGEST (BP) | SPADES N50 | ANIB STRAIN | ANIB (%) |
|---|---|---|---|---|---|---|---|
| **SAMN11961934** | $MD8_{HG}$ | Rumen | 920 | 868,373 | 157,167 | VPI-5482 | 98.03 |
| **SAMN11961935** | $MD11_{MG}$ | Rumen | 62 | 589,645 | 236,443 | VPI-5482 | 98.05 |
| **SAMN11961936** | $MD13_{HG}$ | Rumen | 61 | 781,280 | 229,189 | KLE1254 | 99.16 |
| **SAMN11961937** | $MD17_{HG}$ | Rumen | 59 | 928,731 | 208,464 | KLE1254 | 99.15 |
| **SAMN11961938** | $MD28_{MG}$ | Feces | 63 | 838,230 | 236,443 | 7330 | 98.32 |
| **SAMN11961939** | $MD33_{MG}$ | Feces | 72 | 759,138 | 150,716 | 7330 | 98.33 |
| **SAMN11961940** | $MD35_{MG}$ | Feces | 70 | 723,774 | 145,695 | 7330 | 98.33 |
| **SAMN11961941** | $MD40_{HG}$ | Feces | 62 | 519,222 | 189,229 | 7330 | 98.34 |
| **SAMN11961942** | $MD51_{HG}$ | Feces | 62 | 868,373 | 229,189 | 7330 | 98.33 |

Number of contigs does not include contigs ≤ 1000 bp.

**S3 Table: Statistical significance of RNAseq TPM expression differences between *Bt*VPI-5482, MD33$_{MG}$, and MD40$_{HG}$ MAN-PUL genes when grown in YM.**

| Strain | | TPM | SEM | *B. theta* | MD33 | MD40 |
|---|---|---|---|---|---|---|
| | BT_2620 | 279 | 39 | | ! | - |
| | BT_2622 | 720 | 96 | | ! | - |
| | BT_2623 | 252 | 30 | | ! | - |
| | BT_2625 | 660 | 106 | | ! | - |
| | BT_2628 | 635 | 52 | | ! | - |
| | BT_2629 | 2454 | 385 | | ! | - |
| | BT_2631 | 398 | 64 | | ! | - |
| | BT_2632 | 1087 | 105 | | ns | - |
| | BT_3773 | 179 | 18 | | ! | ! |
| | BT_3774 | 604 | 44 | | ! | ! |
| | BT_3775 | 1646 | 30 | | ! | ns |
| | BT_3776 | 1832 | 62 | | ns | ns |
| [1]***B. theta*** | BT_3780 | 859 | 16 | | ! | ! |
| | BT_3781 | 1103 | 113 | | ns | ns |
| | BT_3782 | 414 | 18 | | ns | ns |
| | BT_3783 | 440 | 52 | | ! | ns |
| | BT_3784 | 1834 | 154 | | ns | ns |
| | BT_3786 | 2594 | 64 | | ! | ns |
| | BT_3789 | 2600 | 358 | | ! | ! |
| | BT_3791 | 265 | 40 | | ! | ! |
| | BT_3792 | 602 | 45 | | ! | ! |
| | BT_3853 | 20 | 2 | | ! | ns |
| | BT_3855 | 12 | 3 | | ! | ns |
| | BT_3858 | 24 | 4 | | ! | ns |
| | BT_3862 | 21 | 2 | | ! | ns |
| | BT_2620 | 13 | 3 | ! | | - |
| | BT_2622 | 16 | 2 | ! | | - |
| | BT_2623 | 24 | 4 | ! | | - |
| | BT_2625 | 804 | 93 | ! | | - |
| | BT_2628 | 183 | 40 | ! | | - |
| | BT_2629 | 2289 | 219 | ! | | - |
| | BT_2631 | 877 | 46 | ! | | - |
| **MD33$_{MG}$** | BT_2632 | 1045 | 300 | ns | | - |
| | BT_3773 | 1201 | 134 | ! | | ns |
| | BT_3774 | 981 | 53 | ! | | ! |
| | BT_3775 | 941 | 157 | ! | | ns |
| | BT_3776 | 1370 | 38 | ns | | ns |
| | BT_3780 | 1635 | 142 | ! | | ! |
| | BT_3781 | 1012 | 163 | ns | | ns |
| | BT_3782 | 1566 | 144 | ns | | ! |

| | | | | | | |
|---|---|---|---|---|---|---|
| | BT_3783 | 1533 | 75 | ! | | ! |
| | BT_3784 | 1301 | 257 | ns | | ns |
| | BT_3786 | 1905 | 146 | ! | | ns |
| | BT_3789 | 132 | 18 | ! | | ! |
| | BT_3791 | 352 | 43 | ! | | ! |
| | BT_3792 | 279 | 49 | ! | | ! |
| | BT_3853 | 449 | 8 | ! | | ! |
| | BT_3855 | 1394 | 45 | ! | | ! |
| | BT_3858 | 826 | 102 | ! | | ! |
| | BT_3862 | 326 | 23 | ! | | ! |
| | BT_3773 | 1261 | 99 | ! | ns | |
| | BT_3774 | 892 | 83 | ! | ! | |
| | BT_3775 | 655 | 22 | ns | ns | |
| | BT_3776 | 3754 | 533 | ns | ns | |
| | BT_3780 | 1677 | 104 | ! | ! | |
| | BT_3781 | 30 | 6 | ns | ns | |
| | BT_3782 | 66 | 17 | ns | ! | |
| | BT_3783 | 36 | 6 | ns | ! | |
| **MD40$_{HG}$** | BT_3784 | 239 | 20 | ns | ns | |
| | BT_3786 | 870 | 128 | ns | ns | |
| | BT_3789 | 237 | 75 | ! | ! | |
| | BT_3791 | 91 | 12 | ! | ! | |
| | BT_3792 | 357 | 66 | ! | ! | |
| | BT_3853 | 107 | 24 | ns | ! | |
| | BT_3855 | 200 | 20 | ns | ! | |
| | BT_3858 | 645 | 128 | ns | ! | |
| | BT_3862 | 163 | 30 | ns | ! | |

[1]*Bt*VPI-5482.

**S4 Table. Percent identity matrix of the MAN-PUL2 SusC/D/E-like amino acid sequences from isolated strains generated by MUSCLE [76].**

| Strain | MD11$_{MG}$ | MD28$_{MG}$ | MD33$_{MG}$ | MD35$_{MG}$ | MD8$_{HG}$ | MD13$_{HG}$ | MD17$_{HG}$ | MD40$_{HG}$ | MD51$_{HG}$ | *Bt*[1] |
|---|---|---|---|---|---|---|---|---|---|---|
| **SusC** | | | | | | | | | | |
| **MD11$_{MG}$** | 100 | 100 | 100 | 100 | 77 | 77 | 77 | 77 | 77 | 77 |
| **MD28$_{MG}$** | 100 | 100 | 100 | 100 | 77 | 77 | 77 | 77 | 77 | 77 |
| **MD33$_{MG}$** | 100 | 100 | 100 | 100 | 77 | 77 | 77 | 77 | 77 | 77 |
| **MD35$_{MG}$** | 100 | 100 | 100 | 100 | 77 | 77 | 77 | 77 | 77 | 77 |
| **MD8$_{HG}$** | 77 | 77 | 77 | 77 | 100 | 100 | 100 | 100 | 100 | 100 |
| **MD13$_{HG}$** | 77 | 77 | 77 | 77 | 100 | 100 | 100 | 100 | 100 | 100 |
| **MD17$_{HG}$** | 77 | 77 | 77 | 77 | 100 | 100 | 100 | 100 | 100 | 100 |
| **MD40$_{HG}$** | 77 | 77 | 77 | 77 | 100 | 100 | 100 | 100 | 100 | 100 |
| **MD51$_{HG}$** | 77 | 77 | 77 | 77 | 100 | 100 | 100 | 100 | 100 | 100 |
| ***B. theta*[1]** | 77 | 77 | 77 | 77 | 100 | 100 | 100 | 100 | 100 | 100 |
| **SusD** | | | | | | | | | | |
| **MD11$_{MG}$** | 100 | 100 | 100 | 100 | 81 | 81 | 81 | 81 | 81 | 81 |
| **MD28$_{MG}$** | 100 | 100 | 100 | 100 | 81 | 81 | 81 | 81 | 81 | 81 |
| **MD33$_{MG}$** | 100 | 100 | 100 | 100 | 80 | 80 | 80 | 80 | 80 | 80 |
| **MD35$_{MG}$** | 100 | 100 | 100 | 100 | 80 | 80 | 80 | 80 | 80 | 80 |
| **MD8$_{HG}$** | 81 | 81 | 80 | 80 | 100 | 100 | 100 | 100 | 100 | 100 |
| **MD13$_{HG}$** | 81 | 81 | 80 | 80 | 100 | 100 | 100 | 100 | 100 | 100 |
| **MD17$_{HG}$** | 81 | 81 | 80 | 80 | 100 | 100 | 100 | 100 | 100 | 100 |
| **MD40$_{HG}$** | 81 | 81 | 80 | 80 | 100 | 100 | 100 | 100 | 100 | 100 |
| **MD51$_{HG}$** | 81 | 81 | 80 | 80 | 100 | 100 | 100 | 100 | 100 | 100 |
| ***Bt*[1]** | 81 | 81 | 80 | 80 | 100 | 100 | 100 | 100 | 100 | 100 |
| **SusE** | | | | | | | | | | |
| **MD11$_{MG}$** | 100 | 100 | 99 | 99 | 78 | 78 | 78 | 78 | 78 | 78 |
| **MD28$_{MG}$** | 100 | 100 | 99 | 99 | 78 | 78 | 78 | 78 | 78 | 78 |
| **MD33$_{MG}$** | 99 | 99 | 100 | 100 | 78 | 78 | 78 | 78 | 78 | 78 |
| **MD35$_{MG}$** | 99 | 99 | 100 | 100 | 78 | 78 | 78 | 78 | 78 | 78 |
| **MD8$_{HG}$** | 78 | 78 | 78 | 78 | 100 | 100 | 100 | 100 | 100 | 100 |
| **MD13$_{HG}$** | 78 | 78 | 78 | 78 | 100 | 100 | 100 | 100 | 100 | 100 |
| **MD17$_{HG}$** | 78 | 78 | 78 | 78 | 100 | 100 | 100 | 100 | 100 | 100 |
| **MD40$_{HG}$** | 78 | 78 | 78 | 78 | 100 | 100 | 100 | 100 | 100 | 100 |
| **MD51$_{HG}$** | 78 | 78 | 78 | 78 | 100 | 100 | 100 | 100 | 100 | 100 |
| ***Bt*[1]** | 78 | 78 | 78 | 78 | 100 | 100 | 100 | 100 | 100 | 100 |

[1] *Bt* = *Bt*VPI-5482. Grey shading represents conservation ≥100%.

**Supplementary Figure Legends**

**Supplementary Fig. 1 Differential glycan utilization of *Bt* strains.** Growth profiles of *Bt*VPI-5482, *Bt*ΔMAN-PUL1/2/3, and nine rumen isolates grown on 0.5% YM extracted from the cell wall of **(a)** *S. cerevisiae* (N=4) and **(b)** *S. pombe* (N=3). Mean ± standard deviation shown.

**Supplementary Fig. 2 Comparative uptake of FLA-YM by *Bt* strains over time**. **(a)** Single cell measurements from pure cultures of *Bt*VPI-5482, $\mathrm{MD33_{MG}}$, and $\mathrm{MD40_{HG}}$. Grey represents cells incubated with YM-MM. Orange shows cells incubated with unconjugated FLA. Blue indicates cells incubated with FLA-YM for the time noted. Graphical representation of how flow cytometry data was gated, with cells above a certain threshold assigned the FLA-YM positive status. N=10.000. **(b)** Epifluorescence images of $\mathrm{MD33_{MG}}$ (left), and $\mathrm{MD40_{HG}}$ (right) incubated with FLA-YM for 0 min, 5 min, and 60 min. Cells were counter stained with DAPI and Nile Red. Scale bar is 10 μm.

**Supplementary Fig. 3 CAZyme updates to PUL85 in the genomes of the *Bt*$^{\mathbf{Bov}}$ isolates.** Grey triangles represent gene inserts into the *Bt*$^{\mathrm{Bov}}$ PUL85-like region relative to *Bt*VPI-5482 PUL85.

**Supplementary Fig. 4 CAZyme analysis of GH92 and GH76. (a)** Synteny of MAN-PUL1/2/3 and HMNG-PUL of *B. theta* and nine *Bt*$^{\mathrm{Bov}}$ strains. Length of genes are to scale and arrows indicate directionality. GH76s are indicated with coloured triangles, whereas GH92s are indicated with coloured circles. MAN-PUL1 = Green, MAN-PUL2 = Blue, MAN-PUL3 = Yellow, and the HMNG-PUL = Red. **(b)** GH92s and **(c)** GH76s characterized and predicted sequences from *Bt*$^{\mathrm{Bov}}$ genomes generated with SACCHARIS(Jones et al., 2018). Large trees (left) show all characterized GH92 or GH76 enzymes from CAZy and those within the *Bt*VPI-5482 genome, as well as the homologous MAN-PUL enzymes from the *Bt*$^{\mathrm{Bov}}$ isolates. Smaller trees display all the characterized GH92 or GH76 enzymes from CAZy, with all of the predicted GH92/76 enzymes found in the genomes of the *Bt*$^{\mathrm{Bov}}$ strain noted above the tree. Outer ring represents characterized specificities (see legend). Circles (GH92) and stars (GH76) represent sequences from PULs: Green = MAN-PUL1, Blue = MAN-PUL2, Yellow = MAN-PUL3, Red = HMNG PUL, and White = PUL55. Numbers in parenthesis indicate the total number of enzymes within each strain.

**Supplementary Fig. 5 Growth profiles of *Bt*$^{Bov}$ cultures supplemented with GH76 recombinant enzymes.** MD33$_{MG}$ and MD40$_{HG}$ cells cultured in YM-MM supplemented with BSA control protein, *Bt*$^{Bov}$ GH76 from PUL55 (BtGH76-MD40), or GH76 from MAN-PUL2 (GH76-BT3782).

**Supplementary Fig. 6 Expression of MAN-PUL gene products in *Bt*VPI-5482, MD33$_{MG}$, and MD40$_{HG}$. (a)** Expression levels of MAN-PUL1/2/3 and HMNG-PUL genes. Grey represents gene transcripts not present in the corresponding genome. **(b)** Transcript expression of MAN-PUL1/2/3 in *Bt*VPI-5482, MD33$_{MG}$, and MD40$_{HG}$. Bar graph represents $\log_2$ fold-change of CAZymes and SusC/D/E-like proteins in MAN-PUL1/2/3 and HMNG-PUL encoded in the genomes of *Bt*VPI-5482 (white), MD33$_{MG}$ (grey), and MD40$_{HG}$ (black). GH76 enzymes indicated by stars. GH92 enzymes indicated by circles. MAN-PUL1 = green. MAN-PUL2 = blue. MAN-PUL3 = yellow. HMNG-PUL = red.

**Supplementary Fig. 7 Co-localization of epitope tagged SusD membrane protein and FLA-YM staining in *Bt*VPI-5482 cells.** Dot blot showing YM-MM (YM) or mannose (Man) cultures of *Bt*Δtdk Δpul75 and a mutant strain with a C-Myc fused SusD-like protein. SR-SIM of *Bt*Δtdk Δpul75 (Bt – wt) stained with DAPI and FLA-YM. SR-SIM of the C-Myc mutant (Bt – cmyc) showing DyLight antibody signal and co-stained with DAPI and FLA-YM. Scales shown are 2 μm.

**Supplementary Fig. 8 Product profiles of YM utilization in the supernatants of *Bt*$^{Bov}$ isolates.** Chromatograms of hydrolysis products in the supernatants of *Bt*VPI-5482, MD33$_{MG}$, and MD40$_{HG}$ when incubated in FLA-YM for 24 hrs (top) and 72 hrs (bottom).
